# Neural habituation enhances novelty detection: an EEG study of rapidly presented words

**DOI:** 10.1101/862516

**Authors:** Len P. L. Jacob, David E. Huber

**Affiliations:** University of Massachusetts, Amherst

## Abstract

Huber and O’Reilly (2003) proposed that neural habituation aids perceptual processing, separating neural responses to currently viewed objects from recently viewed objects. However, synaptic depression has costs, producing repetition deficits. Prior work confirmed the transition from repetition benefits to deficits with increasing duration of a prime object, but the prediction of enhanced novelty detection was not tested. The current study examined this prediction with a same/different word priming task, using support vector machine (SVM) classification of EEG data, ERP analyses focused on the N400, and dynamic neural network simulations fit to behavioral data to provide a priori predictions of the ERP effects. Subjects made same/different judgements to a response word in relation to an immediately preceding brief target word; prime durations were short (50ms) or long (400ms), and long durations decreased P100/N170 responses to the target word, suggesting that this manipulation increased habituation. Following long duration primes, correct “different” judgments of primed response words increased, evidencing enhanced novelty detection. An SVM classifier predicted trial-by-trial behavior with 66.34% accuracy on held-out data, with greatest predictive power at a time pattern consistent with the N400. The habituation model was augmented with a maintained semantics layer (i.e., working memory) to generate behavior and N400 predictions. A second experiment used response-locked ERPs, confirming the model’s assumption that residual activation in working memory is the basis of novelty decisions. These results support the theory that neural habituation enhances novelty detection, and the model assumption that the N400 reflects updating of semantic information in working memory.

## Introduction

Pyramidal cells exhibit temporary synaptic depression, owing to neurotransmitter depletion, which limits the ability of sending cells to signal receiving cells (Abbott et al. 1997). This reduces post-synaptic activity by an order of magnitude, but many theories of object identification (Riesenhuber and Poggio 1999) do not include this dynamic. Furthermore, these theories do not specify how the visual system resets itself for each new visual input. Huber and O’Reilly (2003) proposed that short-term synaptic depression, which in this context we refer to as neural habituation, exists to solve this temporal parsing problem, allowing unobstructed perception of the current stimulus by suppressing the response of recently identified visual objects; because previously viewed objects are suppressed, any new object is highly salient in comparison. However, if an object is repeated, this suppression may make it difficult to identify that object on its second presentation. Huber and O’Reilly developed an artificial neural network model with synaptic depression to explain such repetition blindness effects. However, the benefits of neural habituation were not previously demonstrated. Using computational modeling, support vector machine (SVM) classification of trial EEG, and ERP analyses of the N400, we report evidence that temporary neural habituation as a result of short-term synaptic depression enhances novelty detection when reading rapidly presented words.

According to the habituation model, brief presentations (less than 100 ms) produce a burst of neural activity that carries over, blending with the next visual input. Longer presentations reduce neural activity (reducing blending), making it difficult for previously active cells to respond to new input. Behavioral evidence of this dynamic in low-level vision is found in the ‘missing dot’ paradigm developed by Di Lollo (1980). Subjects report the position of a single missing dot in an array of 5×5 dots shown in two separate presentations, each displaying 12 dots. Accurate performance requires visual integration between the two displays (i.e., blending) and performance increases with increasing duration of the first display up to 100 ms. However, additional increases in duration reduce performance, termed the ‘inverse duration effect’. Prior work documented similar inverse duration effects for higher-level stimuli, such as words (Huber, 2008) and faces (Rieth & Huber, 2010). However, inverse duration effects are not necessarily a detriment when there is a need to parse visual information between each display, rather than integrate across displays. The current study tested the benefits of decreasing visual persistence in a novelty detection task while examining EEG responses to identify the neural basis of enhanced novelty detection.

In the current study, subjects made same/different judgements to each response word in relation to an immediately preceding target word; neural habituation was manipulated by varying the duration of a prime word presented immediately prior to the target word. For brief primes (50ms) there was confusion (blending) between the prime and a different target word, and performance was near chance when the response word repeated the prime. However, when the prime duration was longer (400ms), there was a large increase in accuracy, reflecting the benefits of enhanced novelty detection. We examined the N400 as a neural marker of novelty detection (Kutas and Federmeier 2011), considering that prior work observed smaller N400s for repeated words (Rugg 1985). Such effects are often termed ‘repetition suppression’, and neuronal fatigue has been proposed as a possible underlying cause for this effect (Grill-Spector et al. 2006). Here, we propose that synaptic depression, as an underlying cause of neuronal fatigue, enhances novelty and indirectly affects the N400, while also providing further evidence that synaptic depression can underlie repetition deficits. Our model stands apart from existing perceptual identification and N400 models due to its ability to predict repetition benefits and deficits alike, and to simultaneously predict other ERP components (P100 and N170).

The habituation model is a general account of perceptual dynamics and beyond its application to other word identification effects (Rieth and Huber 2017; Huber et al. 2008b; Potter et al. 2018; Davelaar et al. 2011), it has been applied to repetition effects with faces (Rieth and Huber 2010), categories (Tian and Huber 2010, 2013), spatial attention (Rieth and Huber 2013), and visual scenes (Irwin et al. 2010), in tasks ranging from episodic recognition (Huber et al. 2008a) to the attentional blink (Rusconi and Huber 2018). However, none of the prior studies examined the relationship between neural habituation and the perceptual decision making process (novelty detection). The current study does so by examining EEG signals to the response word across two experiments.

Prior work with the neural habituation model examined perceptual identification tasks as measured with two-alternative forced choice testing (2AFC). To examine predictions regarding novelty detection, the current study used same/different testing so that the neural response to a single response word could be examined. 2AFC testing can be directly related to same/different testing using signal detection theory (Macmillan and Creelman 2005), under the assumption that 2AFC is a direct comparison process whereas same/different testing is a criterial process. A key component of the current study is a specific proposal regarding the neural basis of the criterial process. This extension of the neural habituation model to same/different testing required a new, post-perceptual (i.e., working memory) layer designed to maintain the identities of recently seen words for comparison to perception of the response word. This extension also required a specific measure used for perceptual decision making. In augmenting the model, we assumed that the participant assesses the extent to which the response word was already active in working memory prior to its appearance as the response word. This residual activation for the response word is compared against a criterial value in deciding that the response word is the ‘same’ (more residual activation than the criterion) or ‘different’ (less residual activation than the criterion) than the target word.

The current experiments sought to test these specific neural assumptions regarding the perceptual decision making process. The first experiment did so with non-speeded decision making (a response delay reduced contamination from motor responses) examining stimulus-locked ERPs. This experiment confirmed key predictions regarding the updating of words in the newly proposed maintained semantics layer. The second experiment did so with speeded decision making using response-locked ERPs to test the assumption that residual activation in the maintained semantics layer is the signal underlying perceptual decision making.

## Experiment 1: Stimulus-Locked ERPs

### Material and Methods

#### Experimental design and behavioral data analysis

Twenty subjects aged 18-35 participated in the study. Every participant provided written informed consent, and all study procedures were approved by the University of Massachusetts Amherst Institutional Review Board. Volunteers either received Psychology course credit or a payment of $12/hour as compensation for participating. Subjects were right-handed native English speakers who were neurologically healthy and possessed normal or corrected-to-normal vision.

The experimental task (Fig. 1) was displayed on a 24” LCD monitor with a 120Hz refresh rate. Visual stimuli were generated using PsychToolbox (Brainard 1997; Kleiner et al. 2007) implemented in MATLAB (version 2015a; MathWorks). Each trial began with a fixation cross of a display duration that was adjusted according to prime duration for that trial (650ms on short prime trials, 300ms on long prime trials) such that there was a fixed time between the start of the trial and the briefly flashed target word. After the fixation cross, a blank screen appeared for 300ms followed by a doubled-up prime word. The prime word was doubled-up to provide a visual difference between prime and target even for conditions where the target was the same word as the prime. The prime word was presented for either a short (50ms) or long (400ms) duration.

**Fig. 1.**
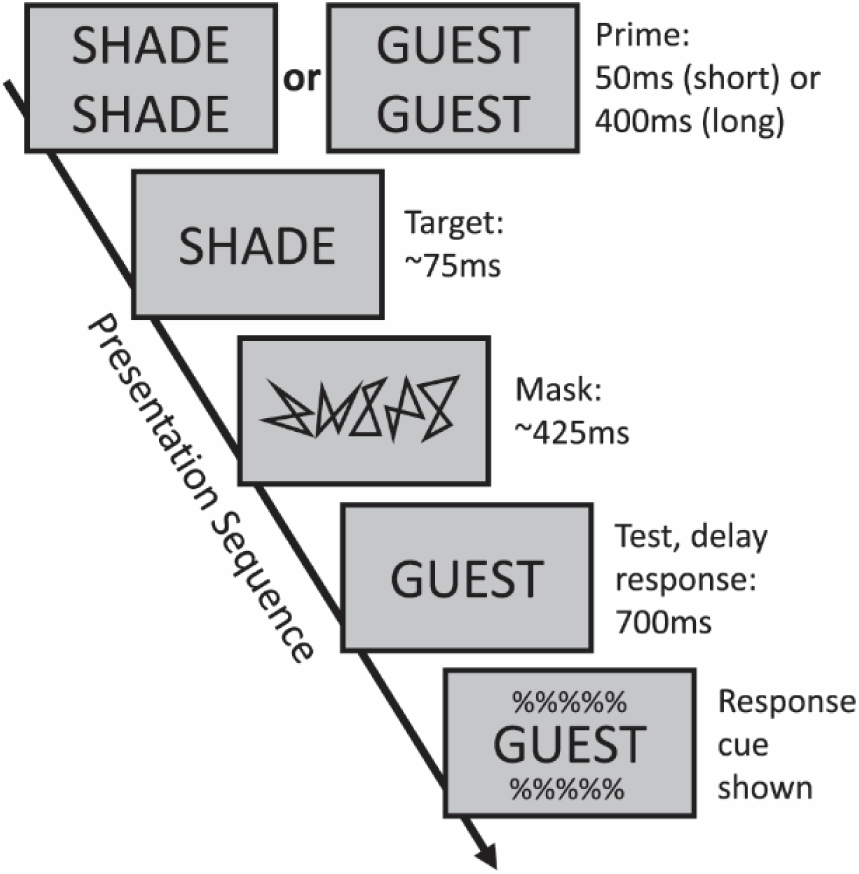
Experimental paradigm. Each trial began with a fixation cross (650ms or 300ms, according to prime duration), followed by a blank screen (300ms) and a doubled-up prime (50ms or 400ms). A target word was then briefly flashed (duration adapted for each subject; average 75ms), followed by a pattern mask (500ms minus target duration) and finally the response display. Subjects were instructed not to press any button until the response cue was shown 700ms later, at which time they made a same/different judgement of the response word in relation to the target word.

A prior ERP study used prime durations of 150ms and 2,000ms with a 2AFC testing version of this task (Huber et al. 2008b). That short duration was chosen because 150ms is long enough to allow explicit identification of the prime; previous work by Huber et al. (2008a) demonstrated that 100ms primes are readable. Thus, the resulting effects could not be attributable to the difference between supra-versus subliminal primes. The short duration in the current experiments was set shorter to maximize positive priming; Huber (2008) ran a study with a range of prime durations from 17ms to 2,000ms, revealing similar positive priming effects at 50ms and 150 ms, but with a larger effect at 50ms. That study also found that both 400ms and 2,000ms produced equally strong negative priming, and so 400ms was chosen for the current experiment to shorten total trial duration. In any event, an analysis of the ERP effects to the target word in the current experiment (not reported here) replicated the target word ERP findings of Huber et al. (2008b) even though each experiment used slightly different short and long prime durations, suggesting that although 50ms may be too brief to allow explicit identification of the prime word, the current results do not reflect the difference between supra-versus subliminal priming.

Immediately following the prime, a target word was briefly flashed; presentation duration was unique to each subject as determined from a block of threshold trials using a staircase adjustment of the target duration to achieve 75% accuracy. Average target word display duration was 75ms. After the target word, a pattern mask was displayed for 500ms minus the target display duration, followed by a single-word response display which remained onscreen. 700ms following initial response word display presentation, a response cue was shown above and below the response word, and subjects had 1.5s to respond; failure to provide a response within that time resulted in a “no response” feedback.

Subjects judged whether the response word was “same” or “different” than the briefly flashed target word. Responses were collected with a custom-made button box with three buttons: a left button, which always corresponded to the response “same”; a right button, which always corresponded to the response “different”; and a center button, which was not used. Subjects held the button box with both hands, pressing the left button with the left thumb, and the right button with the right thumb. To minimize motion artifacts, subjects were instructed to not move or respond until a response cue was shown 700ms after the response word was initially displayed. Feedback was provided immediately after the response, followed by a break to allow blinking/motion that lasted until the subject pressed any button to continue.

There were eight experimental conditions in total, four per prime duration (see Table 1). These conditions were a product of three factors, each with two levels: prime duration (long or short), whether the response word matched the prime word (primed or unprimed), and trial type (same or different). These three factors were used in all behavioral and ERP statistical analyses. It is important to note that “primed or unprimed” can also refer to target word priming (see Table 1), but present analyses focused on the relationship between prime and response words.

**Table 1.**
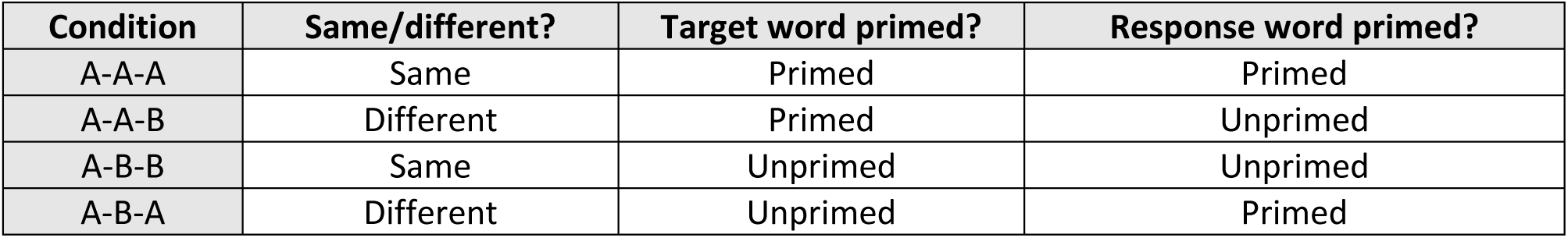
Experimental conditions within each prime duration category.

| Condition | Same/different? | Target word primed? | Response word primed? |
| --- | --- | --- | --- |
| A-A-A | Same | Primed | Primed |
| A-A-B | Different | Primed | Unprimed |
| A-B-B | Same | Unprimed | Unprimed |
| A-B-A | Different | Unprimed | Primed |

In labeling these conditions, we use the letters “A” or “B” to refer to each word in the sequence of events (each trial required no more than two unique words), across the prime, target, and response word presentations. Thus, the same-primed condition is A-A-A (the same word for all three presentations), the same-unprimed condition is A-B-B, the different-primed condition is A-B-A, and the different-unprimed condition is A-A-B.

These 4 conditions stem directly from prior work with 2AFC testing. In prior studies (Huber et al. 2002; Huber and O’Reilly 2003; Huber 2008; Weidemann et al. 2008; Rieth and Huber 2017), the transition from positive to negative priming resulted from comparing a ‘target-primed’ condition to a ‘foil-primed’ condition. In the case of 2AFC testing, this priming referred to which of the two choice words was primed. These 2AFC conditions could be labeled as A-A-A/B (target-primed) versus A-B-A/B (foil-primed) where the final two letters represent the target and foil choice alternatives. The 4 same/different conditions tested here are chosen by breaking each 2AFC condition into two corresponding same/different conditions. Thus, when considering both priming of the target word, and priming of the response word, every condition of the current experiment involved some kind of priming (there was no fully unprimed baseline condition, such as with A-B-C, which is one component of the ‘neither-primed’ 2AFC condition: A-B-B/C).

Words were presented in upper case Arial font size 36, white against a black background. They were randomly drawn without replacement from a pool of 1,087 five-letter words with a minimum written language frequency of 4 per million as defined by Kucera and Francis (1967). Each trial used a unique set of words. Subjects performed 16 practice trials, followed by 80 threshold trials during which target display duration was adjusted every 16 trials in order to achieve the desired 75% accuracy. EEG recording was done during the subsequent 480 experimental trials, which were split into 6 blocks of 80 trials, with a mandatory break of at least 20 seconds between blocks. Every experimental block contained 10 instances of the 8 conditions in a randomly presented order. Between the third and fourth experimental blocks, an electrode impedance check was performed.

#### Habituation model

The habituation model as applied to this paradigm used perceptual dynamics identical to the repetition priming model reported by Huber et al. (2008b) for forced choice testing, with parameter values reported for Experiment 1 of Rieth and Huber (2017). However, same/different testing is different than forced-choice testing, requiring comparison to a response criterion rather than a direct comparison between two alternatives (Macmillan and Creelman 2005). More specifically, the same/different task requires a comparison between the currently displayed response word and recently viewed words to assess the degree of match. The habituation model was thus augmented with a working memory layer representing maintained semantics, with this new layer receiving input from the unchanged base structure that simulates perceptual processing. This new layer was used to generate ERP prediction for the N400 component. We refrained from modifying the base model to highlight the habituation theory’s ability to generalize across subjects and experimental paradigms.

The maintained semantics layer is a completely new addition to the model, with qualitatively different dynamic parameters to enable short-term maintenance of recent semantic representations. It can be contrasted with the episodic familiarity layer employed by Huber et al. (2008a), which captured episodic long-term familiarity by modifying weights related to previously studied words. While the episodic familiarity layer had a slow time constant, the maintained semantics layer updates very rapidly, and does not carry information across trials. The goal of the maintained semantics layer is to hold onto all previous viewed words within the trial, rapidly updating working memory to include the meaning of each word as it appears. To assess whether the response word is ‘same’ or ‘different’ than the target word, the key determinant is how easy it is to update working memory to also include the meaning of the response word. If the response word is already in working memory, it will have a high residual activity in the maintained semantic layer and it will therefore be easy to update working memory to include that word’s identity. However, if that word was not seen previously in the trial, there will be little or no residual activation for that word, making it more difficult to update working memory with that word (a larger N400), suggesting that the correct answer is ‘different’. To achieve this dynamic, there is no inhibition in the maintained semantics layer (no competition between words), the rate of processing is high (large time constant parameter), and maintenance is strong (low leak current and depletion parameters).

The habituation model structure is detailed in Fig. 2. Each node in the model simulates the activity of a large number of neurons with similar inputs and outputs. Model input is determined by all- or-none input (zeros or ones) to the nodes in the retinotopic layer: during the time course of a simulation, while the prime is being presented, the retinotopic prime node receives input of one; when the target is presented, the prime node’s input becomes zero, and the target node’s input becomes one; and so forth.

**Fig. 2.**
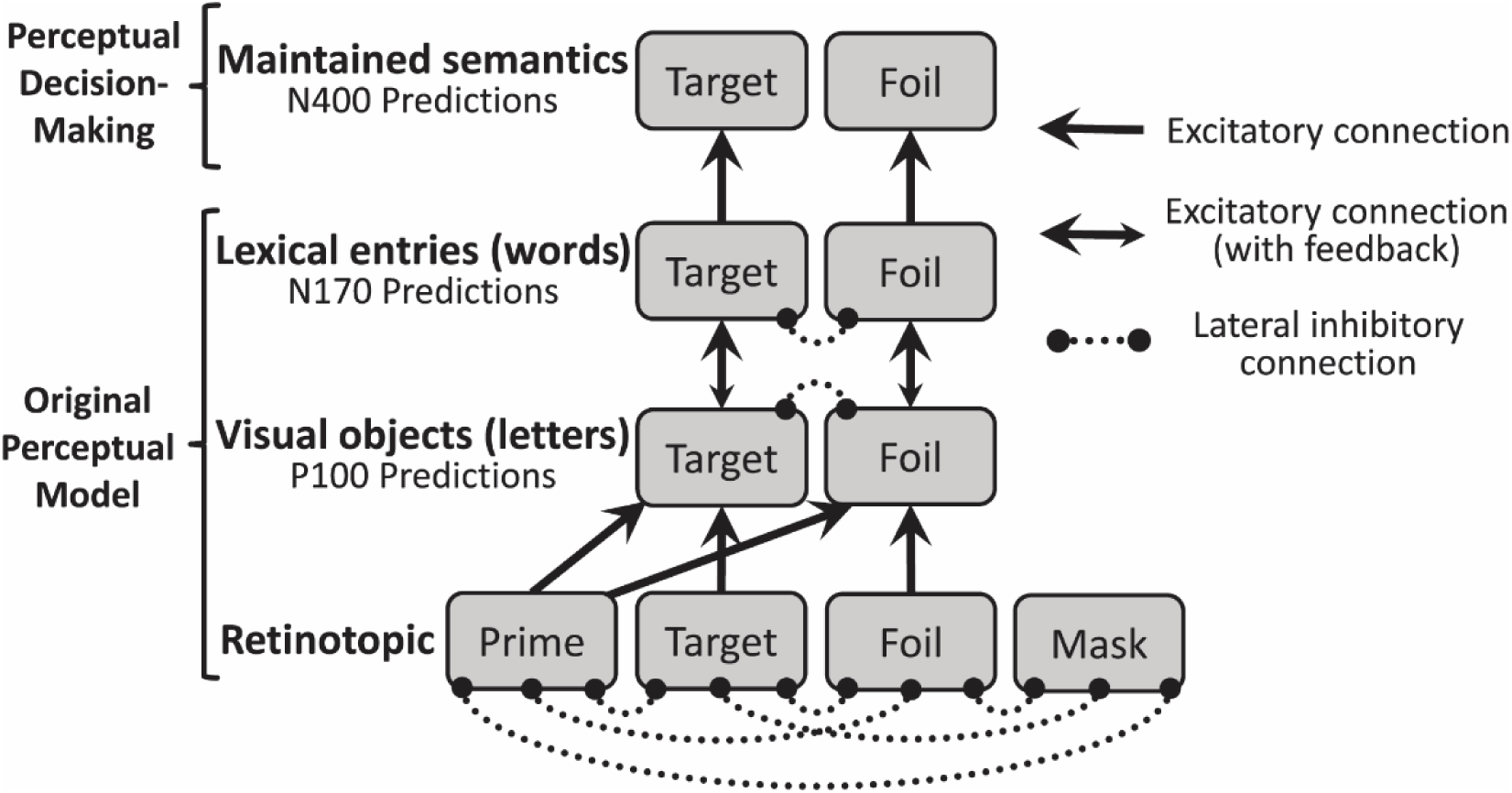
Habituation model structure, implementing the hierarchical nature of visual processing. Each grey rectangle is a node that simulates the activity of a large number of neurons with similar inputs and outputs. The first layer represents retinotopic visual features; the prime node, representing the features of the doubled-up prime word, either connects to the target node (if the prime matches the target) or the foil node (if the prime does not match the target) in the visual objects layer. The retinotopic target node represents the features of the target display, and also the response display, when it is the same as the target. The retinotopic foil node represents the response display when it is different from the target. The perceptual decision making portion is a new addition to the model, capturing the maintenance of each word’s semantic identity in working memory during the course of the trial sequence for comparison with the response word. Updating into this maintained semantics layer is assumed to underlie the N400, and the degree of residual activation for the response word in this layer is assumed to underlie the same/different decision (i.e., more residual activation indicates that this response word was previously seen in the trial, and thus the correct answer is ‘same’).

The activity of the node is captured with two dynamically varying terms, with the product of these determining the signaling that the node can provide to other nodes. The first term is membrane potential (*v*), which is compared to the fixed firing threshold (*Θ*) to determine the probability of an action potential. Because the node implements the activity of many neurons, simulations use this firing rate rather than simulating spiking behavior. However, an action potential does not necessarily produce a post-synaptic response if there are no vesicles available to release, and so the second terms captures the current level of neurotransmitter resources (*a*). Equation 1 is the product of the firing rate and neurotransmitter resources, which is the output of the node.

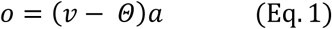

In simulations, these terms are updated every millisecond. At the start of the simulation, output and membrane potential are set to 0, and neurotransmitter resources are set to 1 and the update equations keep these terms between 0 and 1. Membrane potential (*v*) is updated according to Equation 2, which computes Δ*v* for each node *i* in each layer *n*. The first bracketed term corresponds to excitatory inputs, including bottom-up connections from the *n* − 1 layer and top-down feedback (when present) from the *n* + 1 layer, modulated by connection weight *w* (for the bottom-up connection) and feedback strength *F* (for the top-down connection). In the current situation, the weights are set either to 1 or 0 according to whether two nodes are or are not connected. The second bracketed term corresponds to inhibitory inputs, which are a combination of leak (*L*), the natural decay of activation, and lateral inhibition (modulated by inhibition strength *I*), generated by mutual inhibition between the nodes of a layer (and thus affected by their present level of activity). The level of lateral inhibition is the summation of all nodes within the layer, capturing the effect of all-to-all connected inhibitory interneurons, which serve to limit excessive activity. Finally, *S* corresponds to the rate of integration, also unique to each layer.

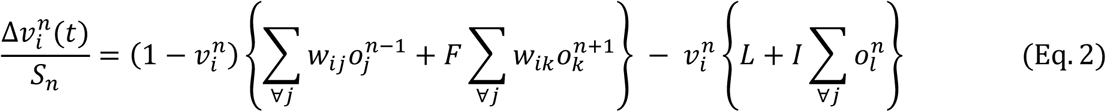

The amount of neurotransmitter resources (*a*) within a node is updated according to Equation 3, which computes Δ*a* as a function of neurotransmitter depletion rate (*D*) and recovery rate (*R*), as well as the node’s output (*o*) and its layer’s rate of integration (*S*).

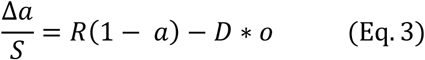

These equations and model structure are exactly the same as in prior publications reporting the simulations with the habituation model. The new component of the model is the maintained semantics layer, which differs from the perceptual layers in the following aspects: it possesses no lateral inhibition (because it needs to maintain the activity of all previously seen words within the trial; lateral inhibition was removed prior to model fitting); the rate of integration is considerably faster (because it needs to rapidly update its contents); the leak value is considerably lower (because it needs to maintain activity); and the depletion value is considerably lower (because it needs to maintain its ability to signal the nature of its contents). Other than the removed lateral inhibition, the parameters were fit to the data. In summary, because the goal of maintained semantics is maintenance and updating rather than identification and temporal parsing, maintained semantics operates quickly, with little synaptic depression and decay, and no inhibition.

The novelty signal used to make behavioral predictions is generated from the maintained semantics layer. This signal determines whether the response word was already in maintained semantics (as would be expected if the response word is a repeat of the target word). If so, there is substantial ‘residual activation’ for the response word in maintained semantics. Residual activation is a measure of short-term familiarity (but not long-term episodic familiarity) and the lack of residual activation reflects novelty. In simulations, residual activation is the minimum output value of the node corresponding to the response word after presentation of the response word (i.e., the low point just before maintained semantics is updated by the response word). If residual activation is above a criterial value (the criterion is a free parameter, but the same criterion is used to simulate performance in all conditions), the model produces a “same” response, but otherwise it produces a “different” response. Supplementary Fig. 1 demonstrates how output and resources in the node corresponding to the response word vary across a trial, and highlights the time in each condition at which this residual activation low-point occurred.

Of note, it might appear that this decision rule should completely fail for non-words. Indeed, Experiment 1 of Rieth and Huber (2017) examined non-word repetition priming with 2AFC testing, finding a similar repetition priming pattern to that of words, with short prime durations producing positive priming and longer durations producing negative priming. The time course of priming was somewhat altered and although a 400ms prime produced negative priming with words, this was the crossover point (neither positive nor negative) for pronounceable non-words, with negative priming failing to appear until the 2,000 prime duration condition. Rieth and Huber assumed that a non-word is reminiscent of several orthographically similar valid words. Therefore, across all lexical entries, the summed activation of non-words may be roughly similar to that of words (e.g., ‘RUDISH’ is reminiscent of ‘RADISH’ and ‘RUSTIC’), but the connection strength between the letters of the non-word and the partially matching lexical entries is weaker. These weaker connections explained the different time course for non-words (i.e., a weaker connection is one that does not lose its synaptic resources as quickly). As applied to the model augmented with a maintained semantics layer, one might assume either that one or more of the reminiscent value words is maintained during the trial or else that participant needs to adopt a different strategy for same/different testing with non-words, such as by maintaining orthographic representations rather than semantic identities. Future work could differentiate between these alternatives through careful manipulation of orthographic similarity in a same/different testing version of this task.

While the model nodes are labeled as “target” and “foil”, the model itself does not use this information to generate its predictions, and cannot internally differentiate between a target and a foil. The reason this labeling system is used is because target and foil words in each trial are both semantically and visually unrelated, dissimilar in meaning and orthography. Therefore, the model structure represents their activation in all layers as two different individual nodes (two different groups of neurons) for simplicity’s sake (since each trial only uses two unique words, a foil and a target, it is unnecessary to model each letter and a complete lexicon). The retinotopic prime is mapped to either target or foil in the visual objects layer depending on which word was being primed (since the prime is doubled-up, it needs its own node in the retinotopic layer, as it does not occupy the exact same screen position as the target word and the response word).

Because the model is deterministic (i.e., every simulation conducted with the same parameters produces exactly the same response), it requires an auxiliary assumption regarding variability to explain different levels of accuracy. For this same/different task, trial-by-trial variability is assumed for the degree of residual activation, with the residual activation from model simulations specifying the mode of this distribution. Because residual activation is bounded between 0 and 1, residual activation variability is captured with a beta distribution. Besides a free parameter for the response criterion for residual activation (*C*), a variability parameter (*N* for noise) specifies the variability of the beta distribution about its mode. In this re-parameterized beta distribution, the *N* parameter can be interpreted as a sample size, dictating the certainty of the residual activation estimate with the stipulation that *N* > 2. The *α* and *β* parameters of the beta distribution are calculated using Equations 4 and 5 and then the probability of responding “different” is a cumulative beta distribution evaluated at *C* with parameters *α* and *β* (the probability of responding “same” is 1 minus this value).

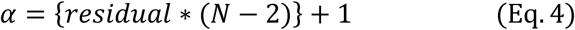

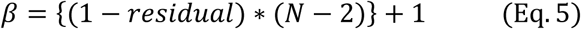

The 5 maintained semantics parameters (*S, L, D, C*, and *N*) were fit to the average observed proportions of “same” and “different” responses across the 8 experimental conditions (16 probabilities reflecting 8 degrees of freedom), minimizing the binomial likelihood ratio test statistic *G*^2^, which is distributed as a χ^2^ (Riefer and Batchelder 1988). The maintained semantics parameter values that best fit the behavioral data were: *S =* 0.3964, *L =* 0.0103, *D =* 0.1036, *C =* 0.0510, and *N =* 38.6155; lateral inhibition (*I*) in this layer was manually set to zero. The parameters inherited from Experiment 1 of Rieth and Huber (2017) were the following: *Θ* = 0.15 and *R* = 0.022 (also used in the maintained semantics layer), *F* = 0.25, *L* = 0.15, *I* = 0.9844, *D* = 0.324; *S* values were the following: 0.0294 (retinotopic), 0.0609 (visual objects), 0.015 (lexical entries). These inherited parameters were obtained by fitting the behavioral accuracy data from a word priming task nearly identical to the one presented here (Rieth & Huber 2017, Experiment 1), except that study examined 5 different prime durations and used 2AFC testing instead of same/different testing. In addition to including a word repetition priming condition, that experiment also tested pronounceable non-words, non-pronounceable non-words, and inverted words, with each producing a slightly different priming time course. As such, that experiment was highly constraining on the parameter values. Because the only major change as compared to that study is the use of same/different testing and augmentation with a maintained semantic layer, the only free parameters in the current study were the 5 new parameters included in these new aspects of the model.

#### ERP Source Modeling

The parameters that best-fit the behavioral data were used to generate predictions for the ERP waveforms. More specifically, predictions for the perceptual dynamics (P100 and N170) were determined from previously reported publications of behavioral data (Rieth and Huber 2017) and the maintained semantics dynamics (N400) were determined from the currently collected same/different behavioral data, and these dynamics were tested by comparing them to the observed ERP data. However, an approximate match to the ERP data requires auxiliary assumptions regarding the manner in which neural sources combine to affect each electrode.

The ERP signal reflects a complex mix of neural sources with the contribution of each source determined not only by the dynamic response of that source, but also by the cortical location of the source (closer to the surface producing a strong localized response versus farther from the surface producing a weaker diffuse response) as well as the orientation of the source (one side of a cortical fold might produce a positive voltage potential while the other side produces a negative voltage potential). Rather than fitting the cortical positions and orientations to the full topographic array of observed ERP data (Berg and Scherg 1994), we limited our analyses to the most relevant electrodes, assuming that different layers of the habituation model contributed in a positive or a negative manner to the data. Thus, for the occipital electrodes used to analyze the perceptual P100 and N170 complex, it was assumed that the visual objects layer contributed a positive response whereas the lexical entries layer contributed a negative response. Similarly, for the central electrodes used to analyze the novelty detection N400, it was assumed that the maintained semantics layer contributed a negative response.

Predictions regarding each source of the ERP data were obtained by summing the output value of all nodes within each layer at each simulated time point (equivalent to a millisecond), and plotting this output over time. Nodes were summed within layers because EEG has low spatial resolution and each neural source reflects a cortical area (e.g., the entire visual word form area rather than separate sources for each word). After obtaining these layer specific activation profiles, they were combined as outlined above in terms of their mathematical sign, with additional specification of how strongly to weight each source and the imposition of delays for each source. In addition, two temporal delay parameters were needed to capture information transfer times not contained in the model (e.g., from the eyes to primary visual cortex, and from word identification in the temporal lobe to working memory in the frontal lobe). In truth these are free parameters, although we chose approximate values that seems to work in general, rather than fitting these anatomical parameters to the ERP data. For completeness and transparency, the layer specific activation profiles are displayed in Supplementary Fig. 2.

These free parameters representing anatomical weighting and delays between different cortical lobes were as follows: The P100 and N170 predictions were created by subtracting three times the lexical entries activation profile from the visual objects activation profile (this choice was made such that lexical entries response was sufficiently strong as to reverse the polarity to produce an N170 after the P100). The retinotopic layer of the model was not included in this summation considering that all words were displayed in the center of the screen; thus, the C1 was not expected to differ between conditions (Gomez Gonzalez et al. 1994). Because retinotopic input in the model reflects early visual cortex, the resulting waveform was temporally delayed by 50ms, reflecting expected retinal and subcortical processing prior to the time when information reaches the visual cortex. Of note, this choice does not change the magnitude of the model’s predictions at all except for changing when the predicted differences between conditions were expected to occur (e.g., whether the P100 effect reaches a peak at 80ms or 120ms). For the N400 predictions, the activation profile of the maintained semantics layer was delayed by 250ms, representing an additional delay of 200ms, reflecting the cortical and sub-cortical connections between perceptual word representations (presumably located in the occipital and temporal lobes) and the updating of maintained semantics within working memory (presumably located in the frontal lobes). In addition, because the central electrodes were topographically adjacent to the occipital P100/N170 electrodes, we assumed some contamination of the central electrodes from ongoing visual objects/lexical entries activity, implemented by adding 0.3 times the predicted P100/N170 waveforms to the value of the predicted N400 waveforms.

Because word reading is not the only neural activity occurring inside a subject’s brain during the course of trial, the model was not expected to explain the ERP waveforms averaged across conditions. A great many other neural sources would need to be included if the goal was to provide a full explanation of the ERP waveforms (e.g., changes in attention, motor preparation, etc.). We assume that these other sources (not included in the model) will be the same across conditions, affecting the ERP waveform averaged across conditions, but not the differences between conditions. The question asked was whether the neural dynamics, as dictated by prior publications (in the case of the P100/N170) and by fitting the dynamics of the maintained semantics layer to the behavioral data (in the case of the N400), could provide a reasonably qualitative account of the ERP differences between conditions.

The ERPs were statistically assessed by averaging across a time window. In theory, one could do the same when comparing the model to data. However, the main reason that a time window is used for statistical analyses of the ERPs is to average over subject differences and trial differences regarding when peak responses occur. For instance, one subject’s peak P100 might occur at 80ms, while another’s occurs at 120ms. Furthermore, even for a given subject, the timing of the peak response will vary from trial to trial. Additionally, time window voltage averages are less affected by noise than peak voltage measures. Because the model does not include subject differences and trial differences, nor does it model voltage variations due to noise, applying a time window to the model is not necessary, and the magnitudes of P100/N170/N400 responses can be directly determined from the simulated time course of events. For predictions of the response word-related P100/N170 complex, we obtained the values for the P100 positive peak and the N170 negative peak; the latter was inverted and summed with the former. For predictions of the response word-related N400, we obtained the values for the negative N400 peak. Finally, we note that the model predictions, as compared to the statistical analyses, needs to be rescaled separately for the P100/N170 complex versus the N400 results, considering that the P100/N170 complex was measured with different electrodes as compared to the N400 (i.e., the Euclidean distances between neural sources and electrodes will be different for each set of analyses).

#### EEG recording and data analysis

Scalp voltages were recorded at a sampling rate of 1000Hz with 0.1-100Hz bandpassing from 64 electrodes (ActiCAP and ActiCHamp; Brain Products). Eye blinks and eye movements measures were collected through auxiliary electrodes (BIP2AUX; Brain products). Individual scalp electrodes were adjusted until each impedance was below 25 kΩ.

EEG and ERP preprocessing and analysis were done using EEGLAB (Delorme and Makeig 2004) and ERPLAB (Lopez-Calderon and Luck 2014) in MATLAB. Continuous EEG data were digitally low-pass filtered at 30Hz, and faulty channels (with kurtosis above 10, or spectrum beyond 5 standard deviations) were rejected and replaced via interpolation. EEG data were re-referenced to an average of all electrodes, epoched, and baseline corrected. Trials with eye blinks or eye movements (as recorded by the auxiliary electrodes and detected by ERPLAB functions) or with large artifacts in the scalp electrodes of interest (see Fig. 3; a threshold of 100µV was used to detect large artifacts) were excluded from all subsequent analyses.

**Fig. 3.**
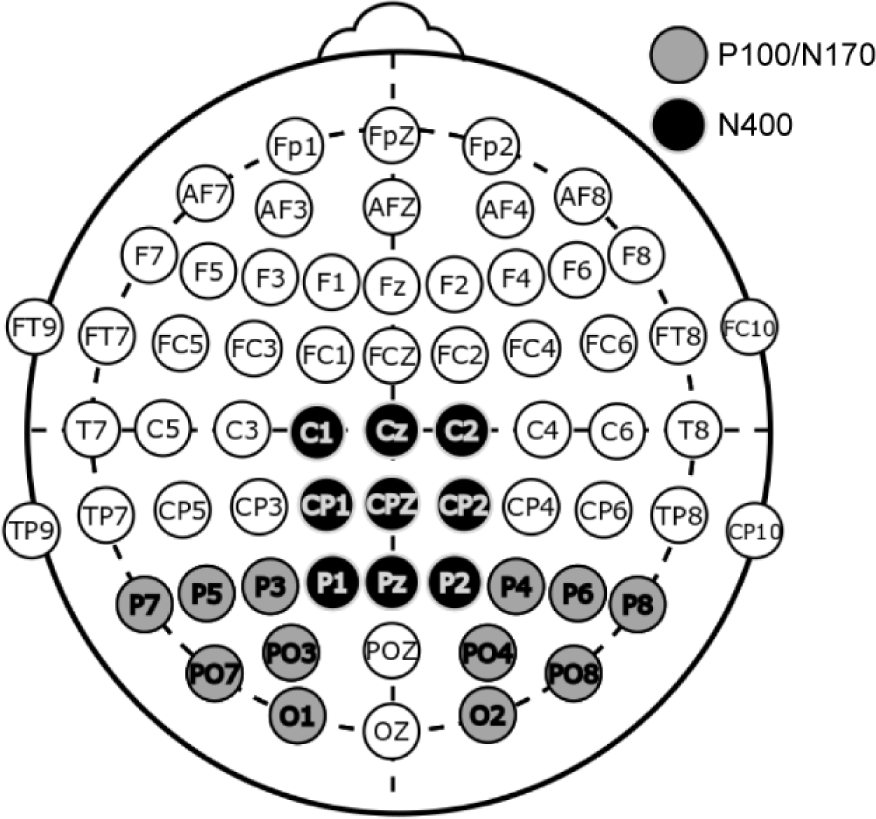
Electrode montage highlighting electrodes used for the P100/N170 complex and the N400 component. Average waveforms were obtained by averaging across the electrodes in each of the highlighted groups.

Given our interest in the N400 to the response word as a measure of novelty detection, a traditional pre-stimulus baseline would confound the analyses by virtue of subtracting relevant ongoing brain activity that is likely to differ between conditions (i.e., the N400 to the target word occurs during the 100ms preceding response word, and if the N400 to the target differ between conditions, this difference in the baseline period would contaminate the analyses). Therefore, we opted for a full-trial baseline as a means of removing electrode drift for the N400 analyses. Epoching was done from −900ms preceding response word presentation (the point in which the doubled-up prime word was presented for trials with long duration primes) up until 700ms after response word presentation (when the response cue was displayed). In this manner, baseline correction utilized the average of the entire epoch. N400 analyses used the average waveform of nine centro-occipital electrodes (Fig. 3). We obtained N400 ERP measures for statistical analysis by averaging the amplitude of the 300-500ms time window following the response word.

We investigated the response word-related P100 and N170 ERP components as a measure of perceptual processing, to further test the habituation model’s predictions regarding perceptual dynamics. Epoching spanned the 100ms preceding response word presentation up until 700ms after response word presentation (the point in which the response cue was displayed). Baseline correction for this epoch used the 100ms preceding response word display (these electrodes do not exhibit the problem of ongoing responses during the baseline period, in contrast to the N400 electrodes). The P100/N170 analyses used the average waveform of twelve occipital electrodes (Fig. 3). ERP measures for statistical analysis were obtained using the signed area method (Luck 2014); positive areas were obtained from a 90-200 ms window, and negative areas from a 140-260 ms window. An additional baseline correction utilizing the data points of 70-280 ms was applied to the waveforms prior to obtaining ERP measures; the goal was to center the P100/N170 complex around a voltage of zero, allowing for more accurate signed area measures. We conducted the ANOVA on the sum of the signed areas of the P100 and N170, with the following justification: in this present paradigm, the stimuli are either repeated words (identical in terms of letters and lexical entries) or unrelated words (different letters and lexical entries), and thus the neural sources of the P100 and N170 should be affected in the same manner by the stimulus, resulting in heavily correlated results. Therefore, we combined these correlated signals to increase statistical reliability.

To create the P100/N170 full-trial plots in Fig. 5 in a manner equivalent to the N400 plots, we epoched the data from the 400ms preceding the target word up until 500ms after target word presentation. This is equivalent to the moment that the prime word is presented (in trials with long duration primes) up until the moment that the response word is presented. We used the time period from −150ms to −50ms in relation to the target word to baseline correct this epoch; this was the 100ms time window with the least amount of contamination from other ERP components. Since two separate baselines were used for the P100/N170 waveforms (as opposed to the N400 waveforms, which used a full-trial average baseline), the natural trial continuity of the EEG signal was broken. To remedy this, the difference between the last time point of the pre-response word P100/N170 epoch and the first time point of the response word P100/N170 epoch was calculated, and then subtracted from each data point in the response word P100/N170 waveform. This resulted in the two epochs lining up to provide a continuous waveform for simpler visualization. These two different baseline corrections are needed because if just one was used, ERPs that were temporally far from that baseline period (e.g., more than 700ms) would have drifted owing to random fluctuations in charge, making comparisons between conditions meaningless. Therefore, our solution is to stitch together these two different baseline corrected epochs, noting that this is in truth a temporal continuum (no time points were removed).

**Fig. 4.**
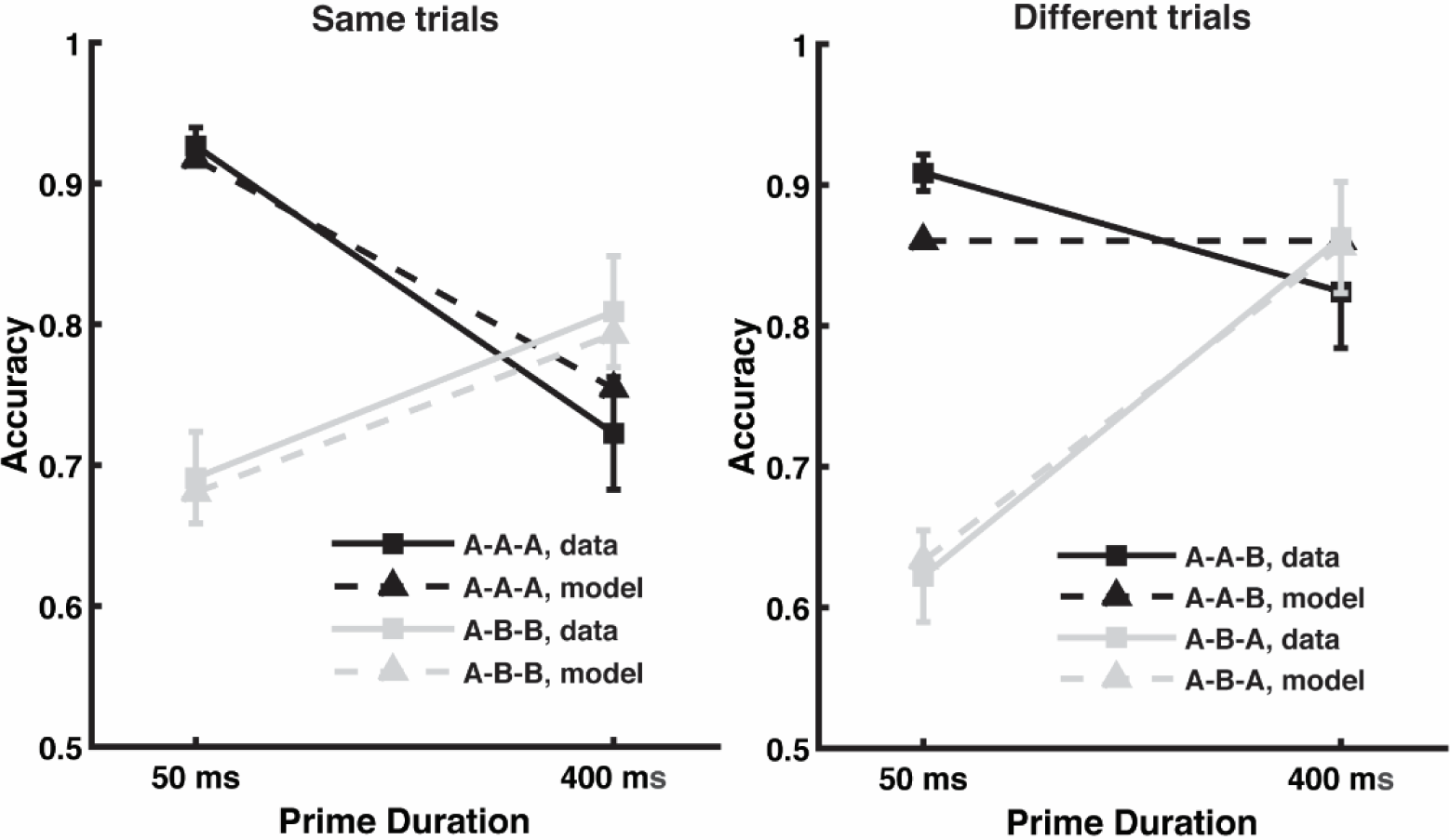
Average subject performance across conditions and equivalent model predictions, with separate graphs for the four conditions where the correct answer was ‘same’ versus the four conditions where the correct answer was ‘different’. The letters “A” and “B” in the legend represent the relationship between the words displayed within a trial, and the order corresponds to prime-target-response. For instance, the A-B-A condition presents one word (word A) as a prime, then a different word (word B) as the briefly flashed target, and then finally the prime word reappears as the response word.

**Fig. 5.**
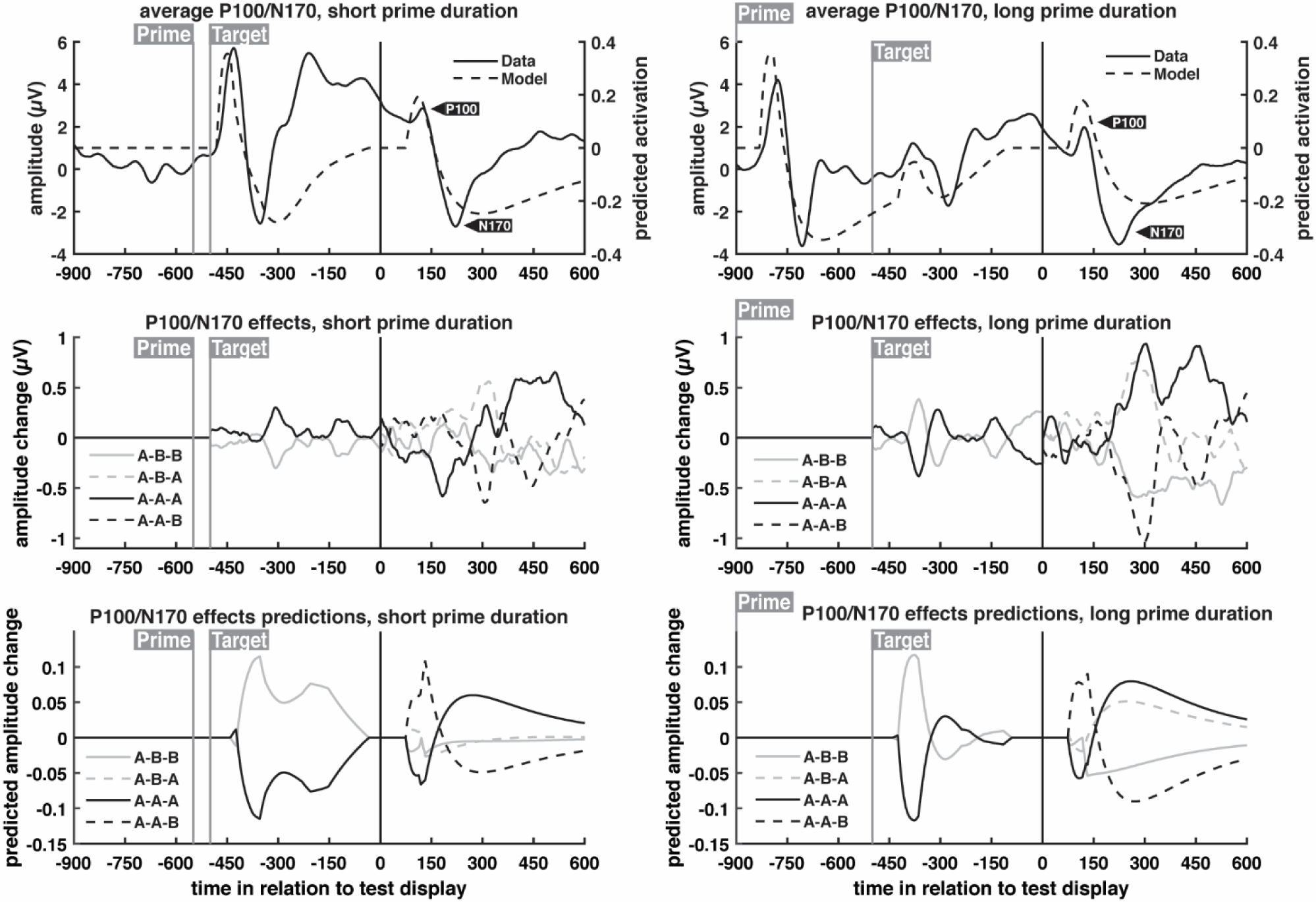
P100/N170 grand average ERP waveforms and model predictions for different prime durations. The first row shows the average of all conditions within each prime duration; subsequent rows show this condition average subtracted from each condition to highlight condition-specific effects. Full trial waveforms are shown, with timing relative to the onset of the response word test display. ERP waveforms were obtained from the average of occipital electrodes (see Fig. 3), while model predictions, based on previously published parameter values, were extracted from the visual objects layer and the lexical entries layer. For layer-specific activation profiles, see Supplementary Fig. 2

#### SVM classification

Using LIBSVM (Chang and Lin 2011) with default parameters, we developed a classifier that could predict choice behavior on a trial-by-trial basis across subjects and conditions to ascertain which electrodes and time points reflected novelty detection. Our a priori hypothesis was that this would correspond to the N400.

The input data consisted of trial EEG epochs spanning the 700ms following response word presentation; these epochs went through the same pre-processing described in the EEG recording and data analysis section above, and were baseline corrected using the average of the entire trial. Only trials in which the subject provided a response were used, regardless of accuracy. The 700ms epochs were divided into 14 windows of 50ms, and the voltage of each window was averaged. This was done independently for each of the 64 electrodes, resulting in a data matrix containing 64 electrodes at each of 14 time periods. This matrix was then reshaped into a vector of 896 features; the classifier, therefore, had no information about the temporal relationship between the time windows or the spatial relationship between the electrodes. Every individual trial for each subject was transformed into a vector of 896 features using the method outlined above, and each vector was normalized with z-scoring.

To parse out variability from random sampling, an SVM classifier was trained and validated 1000 times. Each time, 40 trials for each subject were set aside for validation (5 randomly selected trials from each of the 8 conditions). The remaining trials were used for classifier training (trial number per subject varied, as trials with no response or with EEG artifacts were discarded). For each iteration, classifier performance was validated on all test data, on test data broken down by condition, and on test data broken down by subject; therefore, each iteration produced condition- and subject-specific classification accuracy values, on top of global accuracy. The model received no subject or condition information at any point. These procedures were run 1000 times, and the accuracy values and model weights for each iteration were averaged to provide the final results.

The SVM classifier is non-linear, meaning that the most predictive pattern might involve interactions between different time windows. Therefore, to measure the predictive power of individual time windows in isolation, we also trained and validated the classifier on each time window separately, following the same method above. We then plotted the weights scalp maps of highly-predictive windows to visualize which electrodes were driving the predictions.

## Results

We reiterate that the eight conditions represent combinations of three factors: prime duration (short or long), response word priming (whether the response word matched the prime word: primed or unprimed), and trial type (same or different). To shorten descriptions and allow for easier visualization of the conditions, the letters A and B are used to represent the relationship between the words within a trial. For instance, if a trial began with the prime word GUEST, followed by the target word SHADE, and finally the response word GUEST, it would be an instance of the A-B-A condition (see Table 1).

### Behavioral accuracy results

Behavioral accuracy (Fig. 4) was calculated for each condition across trials in which subjects provided a response. Statistical analysis of accuracy results revealed significant interactions between prime duration and same/different trial type (F(1,19) = 42.07, p < .001), between response word priming and same/different trial type (F(1,19) = 7.93, p = .011), and a three-way interaction between all factors (F(1,19) = 46.59, p < .001). It also revealed a marginal main effect of whether the response word had been primed (F(1,19) = 4.125, p = .0565).

The model captures these results based on the predicted level of residual activation for the response word in the maintained semantics layer as compared to a response criterion. Supplementary Fig. 1 shows the level of residual activation in the model for each of the 8 conditions and the figure caption explains the manner in which residual activation varied across conditions. As seen in Fig. 4, predictions fell within the confidence interval for all conditions other than the A-A-B condition. In this condition, the response word (B) was not seen prior its appearance as the response word, and so the model necessarily predicts that accuracy will be unaffected by prime duration (for both prime durations, there is no residual activation for word B). The observed change in A-A-B accuracy might reflect a situation in which subjects adopt a slightly different response criterion following long duration primes as compared to short duration primes (indeed, a different model fit with two different criteria rather than one was better able to capture the data, although we did not think this level of misfit warranted an additional free parameter). An important caveat to any model fit is the degree of overfitting (too much model flexibility). Demonstrating that the model is highly constrained, Supplementary Fig. 3 shows that the model completely fails to capture these same data, using the same 5 free parameters, if the data are time reversed by labeling the observed data from the 400ms conditions as being the 50ms conditions, and vice versa.

We highlight that the prime can influence behavior in two different ways: by interacting with the target word (making it easier or harder to perceive) and by interacting with the response word (affecting the novelty judgement). We first consider how the prime affects perception of the briefly flashed target word. The main behavioral effect of the prime-target interaction is the performance crossover between A-A (target primed) and A-B (target unprimed) when moving from short to long prime durations. When the prime is presented for a short duration, it blends with the target, improving target identification when they match, but impairing target identification when they do not. When the prime is presented for a long duration, increased habituation from the prime hinders identification of a repeated target, but it also enhances perception of a novel target, owing to less competition from the now habituated prime word.

The largest source of variance in the results is the manner in which the prime interacts with the target (i.e., accuracy for the same trials is largely similar to different trials), but there are important and reliable differences between same trials and different trials (a reliable three-way interaction). The benefits of neural habituation on novelty detection of the response word can be seen when comparing A-B-B and A-B-A trials, both of which entail the same prime-target interactions (both are A-B). Accuracy in the A-B-A condition is at 62% with a 50ms prime (the short prime gives a mistaken sense of familiarity), rising to 86% with a 400ms prime (the familiarity is eliminated, and novelty detection is enhanced). Meanwhile, in the A-B-B condition, accuracy only increases from 69% to 81% (owing to improved target identification). In the model, this occurs because following A-B, the residual activation for a primed response word (A) is reduced with increased habituation, and this reduced residual activation in working memory correctly indicates that the response word A is novel (or at least different than the target word). On the other hand, when comparing A-A-A and A-A-B trials, repetition blindness effects (when moving from short to long prime durations) are much more pronounced on A-A-A trials (worse familiarity detection). In the model, this occurs because following A-A, the residual activation for a primed response word (A) is reduced with increased habituation, and this reduced residual activation in working memory incorrectly suggests that the response word A is novel.

In summary, above and beyond the crossover interaction between prime and target, priming of the response word led to a bias to say ‘different’, with better performance in the A-B-A condition (compared to A-B-B) with increasing prime duration, but worse performance in the A-A-A condition (compared to A-A-B) with increasing prime duration. It is this bias effect that evidences enhanced novelty detection. Next, we analyzed the EEG data to determine the neural correlate of novelty detection and then address whether the neural dynamics correctly predicts the ERP effects.

### ERP results

Average waveforms across all conditions were obtained from both data and model; these are shown in Fig. 5 (for the P100/N170) and Fig. 6 (for the N400). This global average was subtracted from each condition waveform to generate effects waveforms. By plotting the condition averages and the effects separately, it is easier to qualitatively assess the model’s predictions regarding the manner in which neural response were expected to change across the different conditions. Importantly, any additions to model designed to capture the on-average waveforms (averaged across conditions) would not affect the model’s predictions of differences between conditions.

**Fig. 6.**
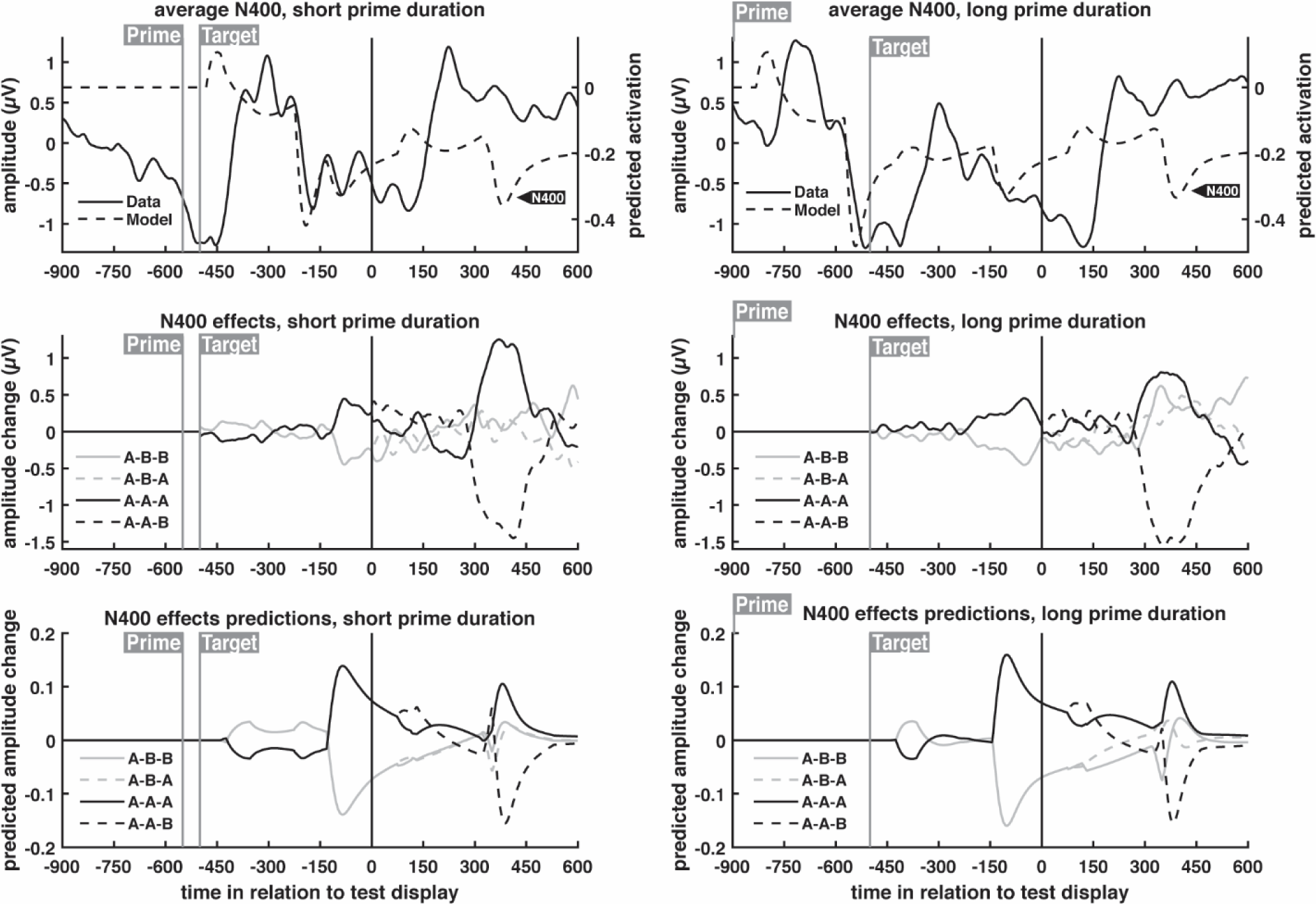
N400 grand average ERP waveforms and model predictions for different prime durations. The first row shows the average of all conditions within each prime duration; subsequent rows show this condition average subtracted from each condition to highlight condition-specific effects. Full trial waveforms are shown, with timing relative to the onset of the response word test display. ERP waveforms were obtained from the average of centro-parietal electrodes (see Fig. 3), while model predictions, based on fits to the behavioral data, were extracted primarily from the maintained semantics layer, with smaller contributions from the visual objects and lexical entries layers.

The occipital electrodes displayed a pronounced P100 and N170 response to the response word, and an ANOVA on the signed areas of the P100 and the N170 revealed an interaction between prime duration and response word priming (Fig. 7; F(1,19) = 12.23, p = 0.002). No other effects were significant. The model was not necessarily expected to capture the results averaged across conditions (because the model only includes word reading areas of the brain), but nevertheless does a reasonable job capturing the major trends. To examine whether the model captured the significant interaction, the absolute value of the predicted P100 and N170 were summed for each of the eight conditions (equivalent to the signed area sum for the real observed data), and then these values were averaged across the ‘same’ and ‘different’ conditions. The model predictions were then rescaled to have the same range as the observed data to place the predictions on the same range as the observed interaction. As seen in the right hand side of Fig. 7, the P100/N170 complex reduced in magnitude for a primed response word, as a function of increasing prime duration, and the model captured this effect, supporting the claim that prime duration increased neural habituation for perceptual representations.

**Fig. 7.**
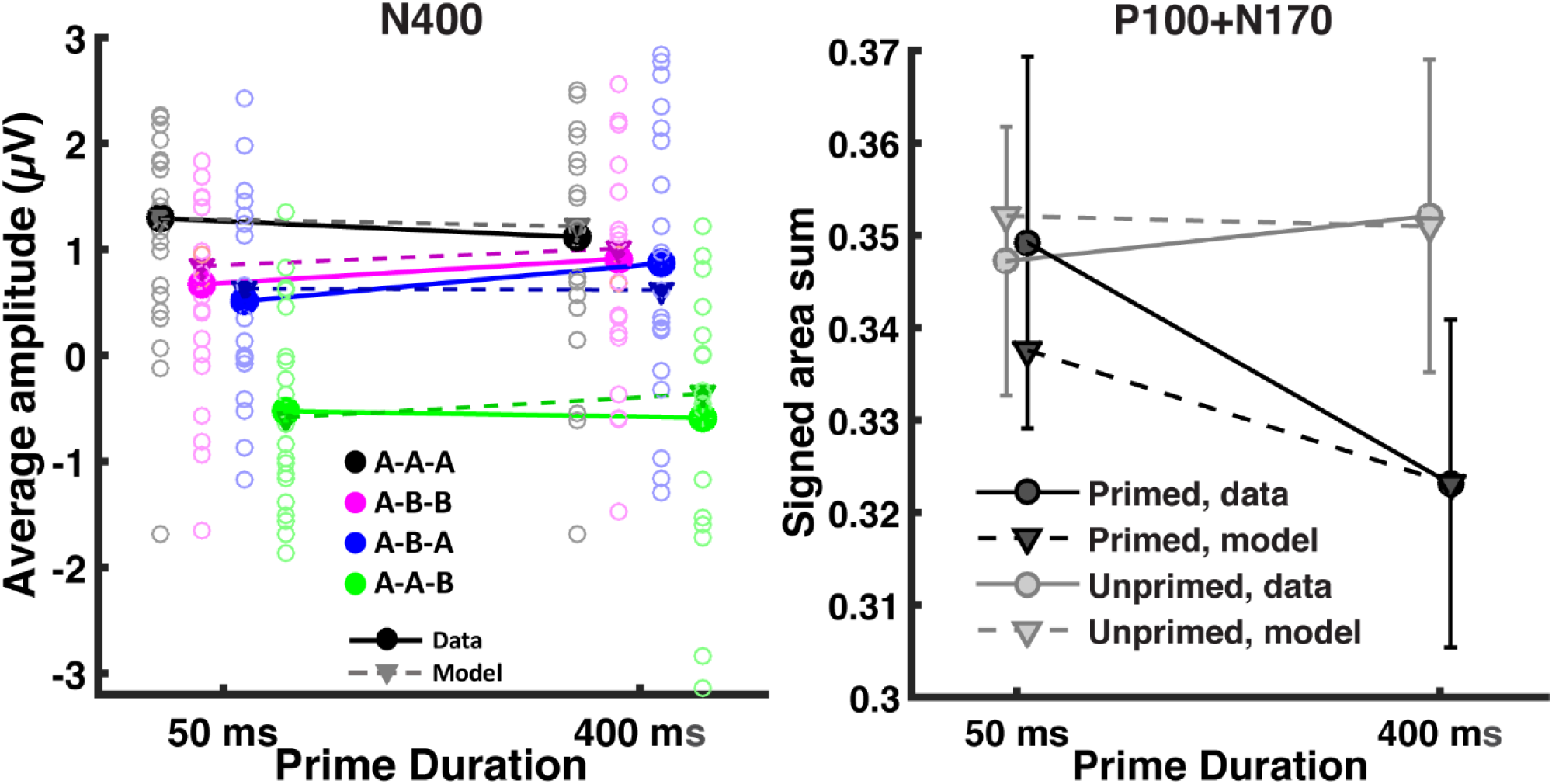
Significant response word ERP results and model predictions. N400 measures were collected from the average amplitude of centro-parietal waveforms between 300ms and 500ms following response word presentation. ‘Primed’ and ‘Unprimed’ refers to whether the response word was or was not a repeat of the prime word. Subject values are represented as circle outlines; some extreme subject values fall outside of the plotted range of values. Response word P100 and N170 measures were collected from the sum of the positive area of the P100 and the negative area of the N170. Cousineau-Morey confidence intervals (Cousineau 2005; Morey 2008) are used to reflect variability in light of large individual differences with this repeated measures design. Model predictions for the P100/N170 were based on previously published parameter values and model predictions for the N400 were based on fits to the behavioral data. To assess whether the model captured differences between conditions, the range of the model’s predictions across the conditions was rescaled to match the range of the observed conditions means.

Next, we consider the manner in which this increased perceptual habituation affected the N400. Centro-parietal electrodes displayed a canonical N400 to both the target word and the response word, characterized by a stronger negativity if the word was seen for the first time in a trial. Although the model did not capture the results averaged across conditions, it closely captured the condition effects.

An ANOVA performed on the average N400 amplitude to the response word (Fig. 7; measured between 300ms and 500ms following presentation) revealed a significant three-way interaction between all factors (F(1,19) = 9.908, p = .005) plus main effects of response word being primed (F(1,19) = 55, p < .001) and same/different trial type (F(1,19) = 40.72, p < .001), along with an interaction between the two (F(1,19) = 9.948, p = 0.005). The magnitude of the N400 ERP is shown in Fig. 7 for all eight conditions, along with equivalent model predictions. The model’s behavior is in terms of output activity (arbitrary units) and so the model predictions were rescaled to have the same range (i.e., the same minimum and maximum values) as the observed data to allow a qualitative comparison between predicted and observed N400 magnitude. While the model’s predictions for the N400 and the P100/N170 share different degrees of contributions from the lexical entries and visual objects layers, rescaling was done independently because the neural sources of the P100/N170 are anatomically distinct from the sources of the N400; therefore, the magnitude detected by a particular electrode will differ (completely distinct electrode groups were chosen for the N400 and the P100/N170).

As seen in the left hand side of Fig. 7, the three-way interaction reflects convergence between the A-A-A and A-B-B conditions with increasing prime duration. The model captured the ordering of the four basic conditions, regardless of prime duration, as well as the three-way interaction that includes prime duration.

### SVM classification to determine which EEG responses predict behavior

The model assumes that residual activation for the response word in working memory underlies behavior; a previously seen word should have more residual activation, making it easier to update working memory to include the response word. This corresponds to a reduced N400 according to the model. To ascertain whether the N400 might be related to behavioral responses, an SVM classifier was trained to predict same/different response from EEG trial data, using one classifier for all subjects and conditions, with an overall accuracy rate of 66.34% (chance = 50.8%—proportion of “different” responses across all subjects and conditions). The classifier was above chance for all subjects; the lowest classification performance when validating on an individual subject was 56.38%. As seen in Fig. 8, the classifier was also above chance for all of the eight conditions.

**Fig. 8.**
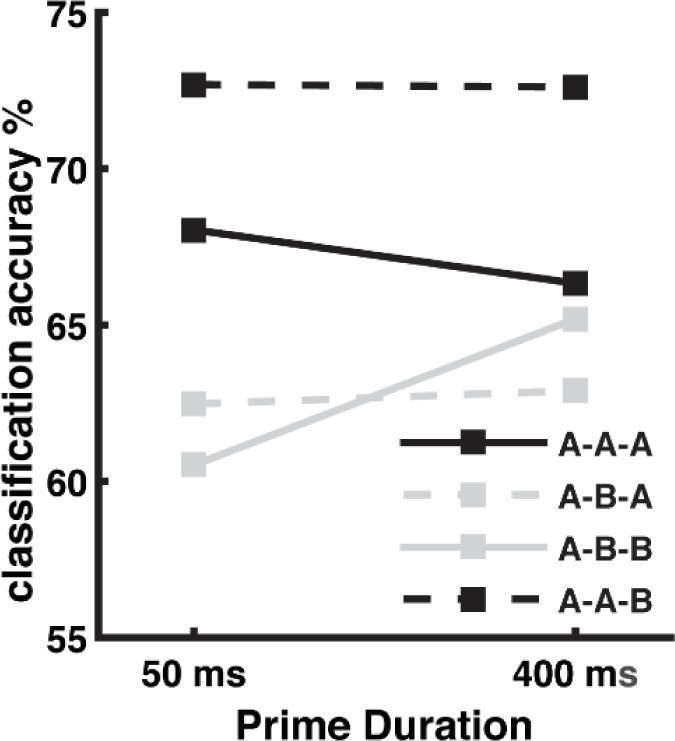
SVM classification accuracy across conditions using held out validation data (see methods for additional details).

To determine which time window is most important for classifier accuracy (i.e., whether N400 amplitude predicts “different” responses on trial-by-trial basis), the classifier was trained and validated on each time window separately; classification accuracy over time is shown in Fig. 9a. Accuracy in the initial time windows is close to chance, until it begins rapidly increasing with the 200-250ms window, finally peaking at 61.14% in the 350-400ms window. Afterwards, it decreases sharply, rising again over the final three time windows, reaching 56.87% in the 650-700ms window, which immediately precedes the cue indicating that the subject can now respond. We hypothesize that the high-accuracy classification value in the 350-400 window reflect the N400, whereas the moderate classification accuracy rise prior to the response cue reflects motor planning.

**Fig. 9.**
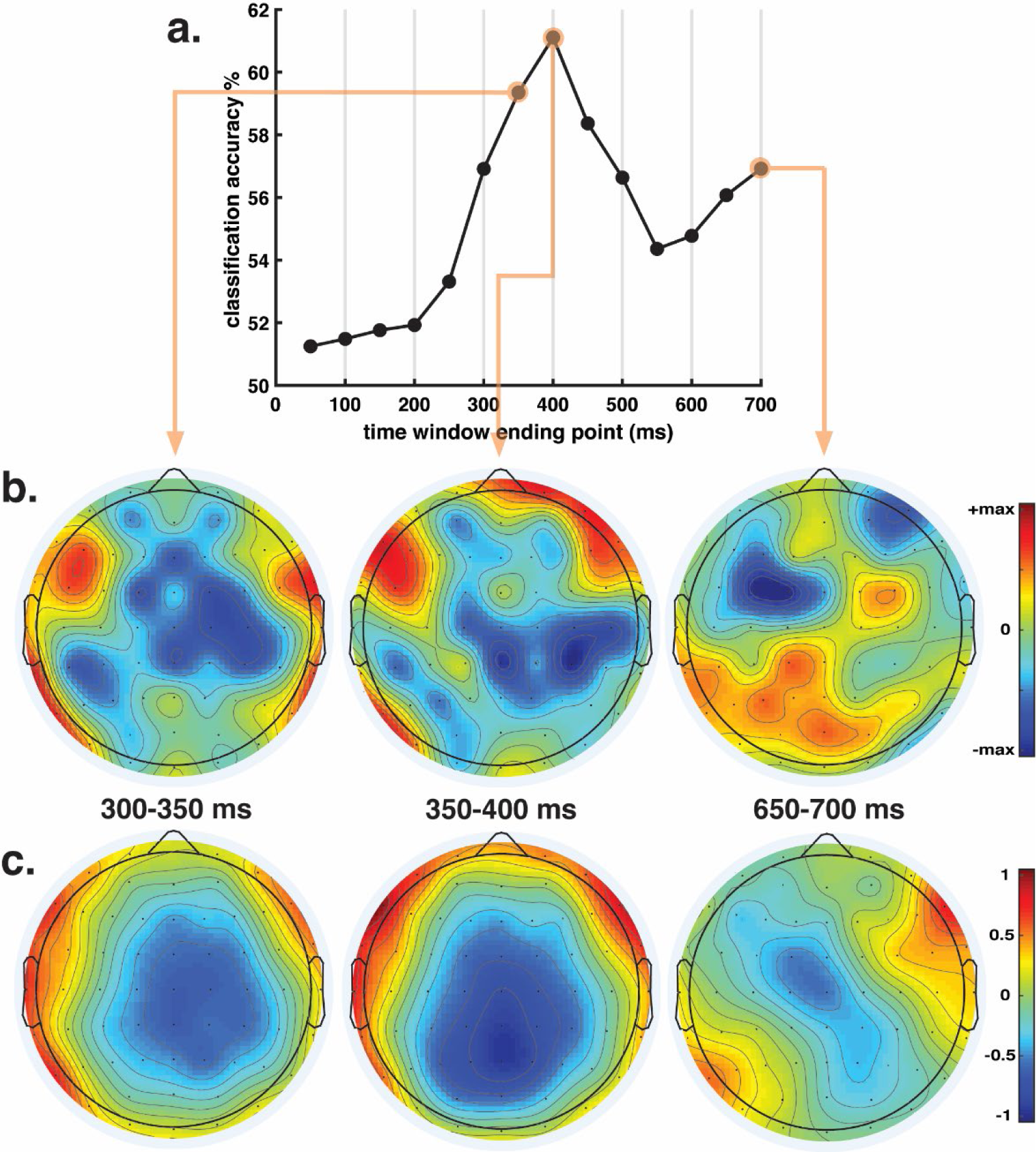
SVM time window analysis. ***a.*** Classifier performance when trained and validated on each individual time window. ***b.*** Scalp plots of the electrode weights from the highlighted windows. The max scale value for 300-350 and 350-400 is 40; the max value for 650-700 is 80. ***c.*** Equivalent grand average ERP scalp plots to the three time windows displayed in b, showing the mean voltage difference (in μV) between trials in which subjects responded “different” minus trials in which they responded “same”.

Interpreting the weight vectors of a classifier is tricky, as significant nonzero weights may be observed at channels that do not reflect the neural process of interest (Haufe et al. 2014). Nevertheless, a rough characterization of the weights is gained by plotting the topography of the weights at each time window of interest (Fig. 9b) next to the observed ERP data (Fig. 9c - grand average voltage difference between trials in which subjects responded “different” minus trials in which subjects responded “same”). The model was coded with “same” responses as −1 and “different” responses as +1; therefore, in the presence of negative weights, more negative voltages predict a “different” response, while more positive voltages predict a “same” response. The opposite is true for positive weights. With this in mind, the 300-350 and 350-400 windows revealed strong negative weights that overlap with the observed topography of the ERP data voltage difference; in the data, trials in which subjects responded “different” displayed stronger central negative voltages between 300-350 and 350-400 than trials in which subjects responded “same”; the model seems to be using this difference (which is, essentially, the N400) by placing strong negative weights in this area (which, paired with more negative voltages, would lead to a “different” prediction). In contrast, the 650-700ms time window revealed a strong asymmetry between the hemispheres consistent with preparation to press one button or the other button (the left button was used to indicate “same” and was pressed by the left thumb whereas the right button was used to indicate “different” and was pressed by the right thumb). This asymmetry was present both in the weights topography and the observed ERP data.

## Experiment 2: Response-Locked ERPs

The behavioral data from Experiment 1 indicated that novelty detection was enhanced with increasing prime duration and the P100/N170 results suggest that neural habituation was the cause of this effect, with the magnitude of the P100/N170 response to a primed response word decreasing with increasing prime duration. Prime duration also affected the N400, with the two “same” conditions (A-A-A and A-B-B) producing a similar magnitude N400 following a long duration prime. Application of the model to the behavioral results assumed that that key measure of performance was not summed maintained semantics activation, which is the signal used to model the N400, but rather just the component of maintained semantics activation that is unique to the response word. Better measurement of this response word response is addressed in Experiment 2, which replicated Experiment 1, but with a response-locked design rather than stimulus-locked.

Experiment 2 was identical to Experiment 1 except that there was no response cue and this experiment asked subjects to respond quickly once the response word appeared. The goal of this experiment was to test the model’s assumption regarding maintained semantics residual activation as the novelty detection variable underlying the decision process. More specifically, because the timing of the response to the response word is likely to vary across trials, this response-locked paradigm should temporally align trials based on the subject’s decision to respond, allowing a cleaner measure of the neural activity leading up to the novelty detection decision.

A different cohort of 20 subjects was used in this experiment; other than the instruction to provide speeded responses, every aspect of experiment design, EEG acquisition, and EEG pre-processing was the same. EEG epochs spanned the 600ms preceding the moment a response was made; baseline correction utilized the average of the full epoch. Mean waveforms were obtained by averaging across the same N400 centro-parietal electrodes (see Fig. 3). ERP measures for statistical analysis were obtained from the average voltage of the 100ms preceding the response.

## Results

EEG epochs were locked to the moment the response was made, and averaged to produce ERP waveforms. We examined the mean average voltage for the 100ms preceding the response to isolate response related activity. Voltages were obtained from the same set of centro-parietal electrodes used for the N400 analyses of the first experiment (see Fig. 3). Mean voltages and model predictions were z-scored within each prime duration to isolate the interaction between prime duration and condition separate from any main effect of prime duration (Fig. 10).

**Fig. 10.**
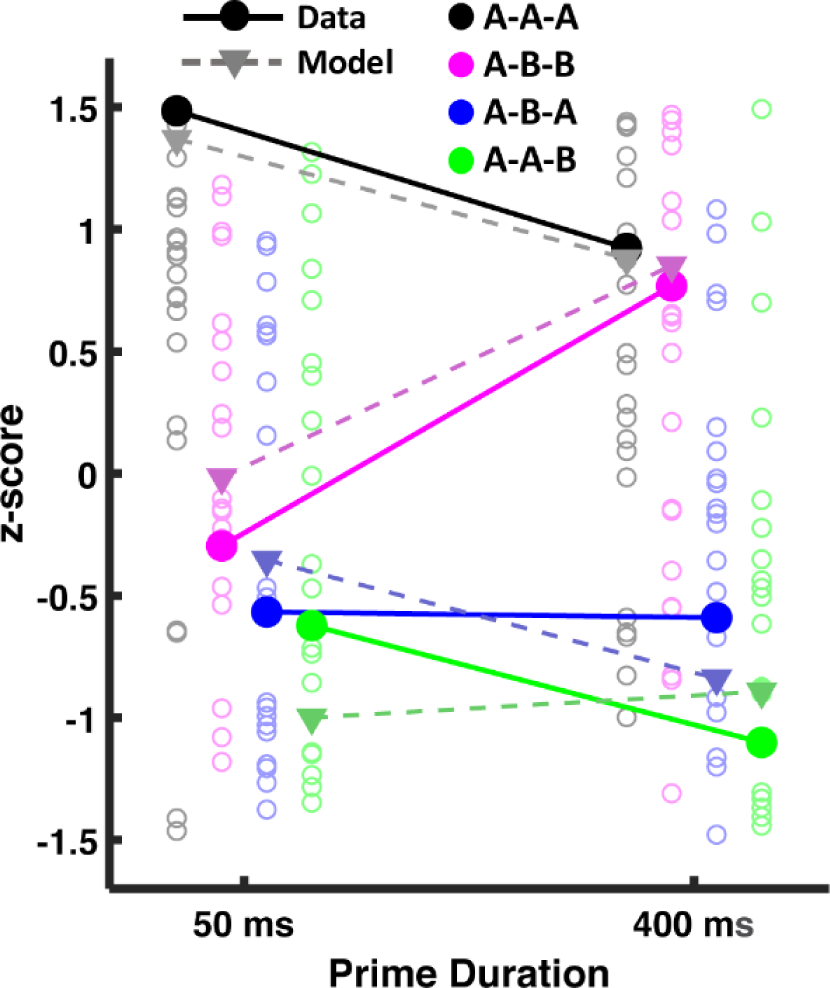
Pre-response ERP measures were obtained from the mean voltage of the 100ms preceding the response for the N400 electrodes. Subject values are represented as circle outlines. Model predictions correspond to the variable assumed to underlie the same/different decision, which is the lowest activation value of the response word’s node in maintained semantics (i.e., residual activation for that word), after presentation of the response word. As seen in the figure, increasing prime duration differentiated same trials (A-A-A and A-B-B) from different trials (A-B-A and A-A-B). All values were z-scored separately (for average data results, for each subject, and for the model) within each prime duration.

Model predictions in Fig. 7 for the N400 reflected the maximum of the summed activation of the maintained semantics layer. However, for response locked results, the neural data reflect signals preceding the decision, rather than signals locked to the onset of the response word. We assumed that response-locking isolated signals unique to the decision process (rather than the summed response across maintained semantics), and so the measure used to explain behavior (i.e., residual activation for the response word) was directly used in making predictions for the response-locked data. As seen in the figure, pre-response voltages displayed the same ordering of conditions as the predicted maintained semantics residual activation values. An ANOVA performed on pre-response mean voltages revealed a significant main effect of prime duration (F(1,19) = 14.9, p = 0.001), with increased voltages on trials with long duration primes, and of same/different trial type (F(1,19) = 20.04, p < .001). It also revealed a marginal three-way interaction between all three factors (F(1,19) = 3.39, p = 0.08) in the form of convergence between primed and unprimed same trials (A-A-A and A-B-B) whereas prime duration had little effect on the two kinds of different trials (A-B-A and A-A-B). According to the habituation model, the convergence between the ‘same’ conditions reflects increased neural habituation in response to the prime word. In the A-A-A condition, this increased habituation produces weaker lingering activation to the target word A, and so the residual activation of this word in maintained semantics at the time of response word presentation is lower. On the other hand, in the A-B-B condition, the increased habituation to the prime word A allows the representation of the target word B to reach higher levels of activation due to less lateral inhibition from A, resulting in greater lingering activation for the target word B.

## Discussion

According to the habituation model, neural habituation as a result of resource depletion is beneficial, clearing the way for rapid identification of new visual objects with minimal interference from recently viewed objects. Thus, short-term synaptic depression enhances novelty detection. However, repetition blindness is a necessary side effect of neural habituation. Habituation is assumed to exist at all levels of perceptual processing, from low-level perception of lines, motions, and colors (e.g., color aftereffects) to high-level perception of letters and lexical entries processing (e.g., negative word priming). A prior ERP study found evidence of neural habituation in a word priming paradigm that produced negative priming, and the habituation model successfully predicted P100 and N170 repetition suppression effects in response to the target word (Huber et al. 2008b). However, that study was not able to measure the separate neural responses to response words that did or did not match the target word because the response display included both a matching and a mismatching word for a forced choice decision. Consequently, the prior study did not assess whether neural habituation supported enhanced novelty detection (e.g., an enhanced response to a response word that differed from the target). The current experiments tested this hypothesis by using same/different testing rather than forced choice responding. This allowed examination of neural responses to a single response word following presentation of the prime and target words, with the results supporting the hypothesis that resource depletion in response to a prime (reduced P100/N170 with increasing prime duration) enhances novelty detection.

Enhanced novelty detection was evidenced both behavioral and neurally. Above and beyond interactions between the prime word and target word (which were documented in a number of previous publications with 2AFC testing), there was an increased bias to respond “different” for response words that matched the prime, as a function of increasing prime duration. This resulted in larger improvements in the A-B-A condition (as compared to the A-B-B condition) but also larger deficits in the A-A-A condition (as compared to the A-A-B condition) as a function of increasing prime duration. Experiment 1 examined stimulus-locked ERPs, finding that perceptual responses (P100/N170) to primed response words decreased with increasing prime duration, providing evidence of perceptual habituation. More importantly, there was a complex three-way interaction for the N400 as a function of prime duration, priming status, and same/different status, which corresponded to qualitative predictions based on the summed working memory activity in the neural network model based on parameters that best fit the behavior data. To isolate the signal unique to the response word in the decision process, Experiment 2 was identical to Experiment 1, but examined speeded response-locked ERPs, confirming the prediction that with increasing prime duration, the neural signal for the same conditions became better separated from the different conditions.

An increased N400 can occur in response to a word that violates semantic expectations—e.g., “I like my coffee with cream and SOCKS” (Lau et al. 2008)—and we anticipated that a different response word (as opposed to a response word that was the same as the target word) would produce a larger N400 considering that the target word and response word were semantically unrelated. To test whether this semantic novelty was the basis of behavior, we used an SVM classifier to predict trial-by-trial behavioral responses from EEG, finding the greatest predictive power around 400 ms after the presentation of the response word, with a topographic pattern that is consistent with the N400. In modeling the N400 and its relation to behavior, we assumed that the meaning of each word is encoded into maintained semantics, with the N400 reflecting this encoding. With this assumption, a word that is semantically expected is a word that is partially active in maintained semantics, whereas the encoding of an unexpected word into maintained semantics requires a more substantial update. Thus, an effective measure of whether the response word differs from the immediately preceding target word is the degree of residual activation in working memory for that word.

To implement these theoretical assumptions, the maintained semantics layer was assumed to exhibit relatively less synaptic depression and no inhibition, allowing it to maintain the identity of previously presented words for longer periods of time than the perceptual layers. The dynamics of the perceptual layers were specified by previously published parameter values and the dynamics of the maintained semantics layer were adjusted to fit the behavioral data. These parameter values were then fixed to generate a priori predictions for the full-trial ERP waveforms for all eight of the conditions representing combinations of prime duration, primed/unprimed, and same/different response words. The examined electrodes captured the P100, N170, and N400 responses, corresponding to visual objects, lexical entries, and maintained semantics processing in the model. The model only included these three components and as such could not provide a full quantitative account of all neural behavior, but nonetheless the model produced a qualitative account of the ERP differences between conditions. The success of this account not only supports the proposal that synaptic depression enhances novelty detection, but also supports the auxiliary assumption that the N400 reflects the process of loading new content into maintained semantics, explaining why the N400 is smaller for an expected word (i.e., semantic content that is already active based on the preceding sentential context).

Providing support for the conclusion that semantic novelty is the basis of performance in this word identification task, the SVM classifier was able to predict choice behavior across subjects and conditions, and the time window with highest predictive power was between 350 and 400ms after presentation of the response word, with a weight topography consistent with the N400. Demonstrating that the N400 is a robust predictor of behavior, classifier accuracy was 66.34%, and was above chance for all subjects and conditions. The manner in which classifier accuracy differed across conditions provided further support for this conclusion: the classifier achieved its best performance for A-A-B trials, which produced the largest N400; second best performance for A-A-A trials, which produced the weakest N400; and worst performance for the A-B-A and A-B-B conditions, which produced intermediate N400 effects.

Despite the success of the SVM classifier for the stimulus-locked experiment, the neural habituation model assumes that residual activation in working memory is the neural correlate of the decision process, rather than the magnitude of the N400. In general, more residual activation should result in a smaller N400 and less residual activation should result in a larger N400, but there may be subtle differences between these measures. To better isolate the hypothesized decision-related response, we performed a response-locked version of the experiment, examining N400 electrodes during the time period immediately preceding the response. This response-locked response was affected by prime duration, exhibiting the same ordering of conditions as the residual activation values from the model.

Our approach in using a layered dynamic neural network to make sense of ERP data highlights the potential limitations of treating separate ERP components as separate processes (see Anderson et al. (2016), for a more statistical approach for tackling this problem based on trial-by-trial variability). For instance, in separate studies, the P100 is examined as a marker of mid-level visual processing and visual attention (Foxe and Simpson 2002; Di Russo et al. 2002), the N170 is examined as a marker of face processing or expertise (Kappenman et al. 2012), and the N400 is examined as a marker of semantic novelty (Kutas and Federmeier 2011). In contrast, we assume that all layers of processing contribute to all of these ERP waveforms, but with each ERP more strongly driven by a specific layer, with higher layers achieving maximal activation at longer delays after stimulus presentation. Thus, in applying the habituation model to the ERP data, visual objects identification is the primary determinant of the P100, lexical identification is the primary determinant of the N170, and the updating of maintained semantics is the primary determinant of the N400, and yet there is contamination of the ERP responses from all layers. Untangling this contamination requires a formal model to specify the dynamic time course of each layer and our approach used the behavioral data to specify this time course. This can be contrasted with source modeling algorithms that impose no constraints on the time course of the neural sources (Baillet et al. 2001).

The habituation model is primarily a perceptual identification model, rather than an N400 model; however, it provides a novel and easily-generalizable explanation for the neural basis of this component that is consistent with prior results. In some respects, prior N400 models are similar to the maintained semantics layer in the current model (Frank et al. 2015; Cheyette and Plaut 2017; Rabovsky et al. 2018; Laszlo and Plaut 2012; Brouwer et al. 2017; Laszlo and Armstrong 2014; Rabovsky and McRae 2014), but these prior models are not well-suited to explain repetition priming effects and the shift from positive to negative priming with increasing prime duration. This extension of the habituation model assumes that perceptual dynamics, and more specifically synaptic depression, is the root cause of this shift, and a complete account of the N400 should incorporate the manner in which perceptual dynamics interact with semantic novelty to produce an N400 effect.

This study is a part of a larger investigation into the proposal that short-term synaptic depression is adaptive, serving to temporally parse the stream of constantly changing perceptual inputs. Many paradigms reveal behavioral effects that rise and fall with increasing presentations and delays, including the inverse duration effect (Di Lollo and Dixon 1992; Di Lollo and Bischof 1995), priming of depth perception (Long et al. 1992), repetition word priming (Burt et al. 2014), semantic word priming (Rieth and Huber 2017), face priming (Webster and MacLin 1999), response priming (Eimer and Schlaghecken 2003; Lleras and Enns 2004), inhibition of return (Posner and Cohen 1984), the psychological refractory period (Pashler 1994), and the attentional blink (Chun and Potter 1995). The time course of these effects is remarkably similar, and as reviewed in the introduction, the habituation model has been applied to many of these “cognitive aftereffects” and explained them. According to the habituation model, these effects reflect a deficit when something repeats for sufficiently long (i.e., a repeated dot, depth plane, word, face, response, location, response selection, or categorically defined target). However, this study is the first true test of the model’s proposal that synaptic depression is beneficial, serving to accentuate responses to a novel stimulus.

## Acknowledgments

We thank Anushree Mehta for her work on an earlier version of the SVM classifier, and Christoph Weidemann for his suggestions towards improving our classifier analyses.

---

**Supplementary Fig. 1.**
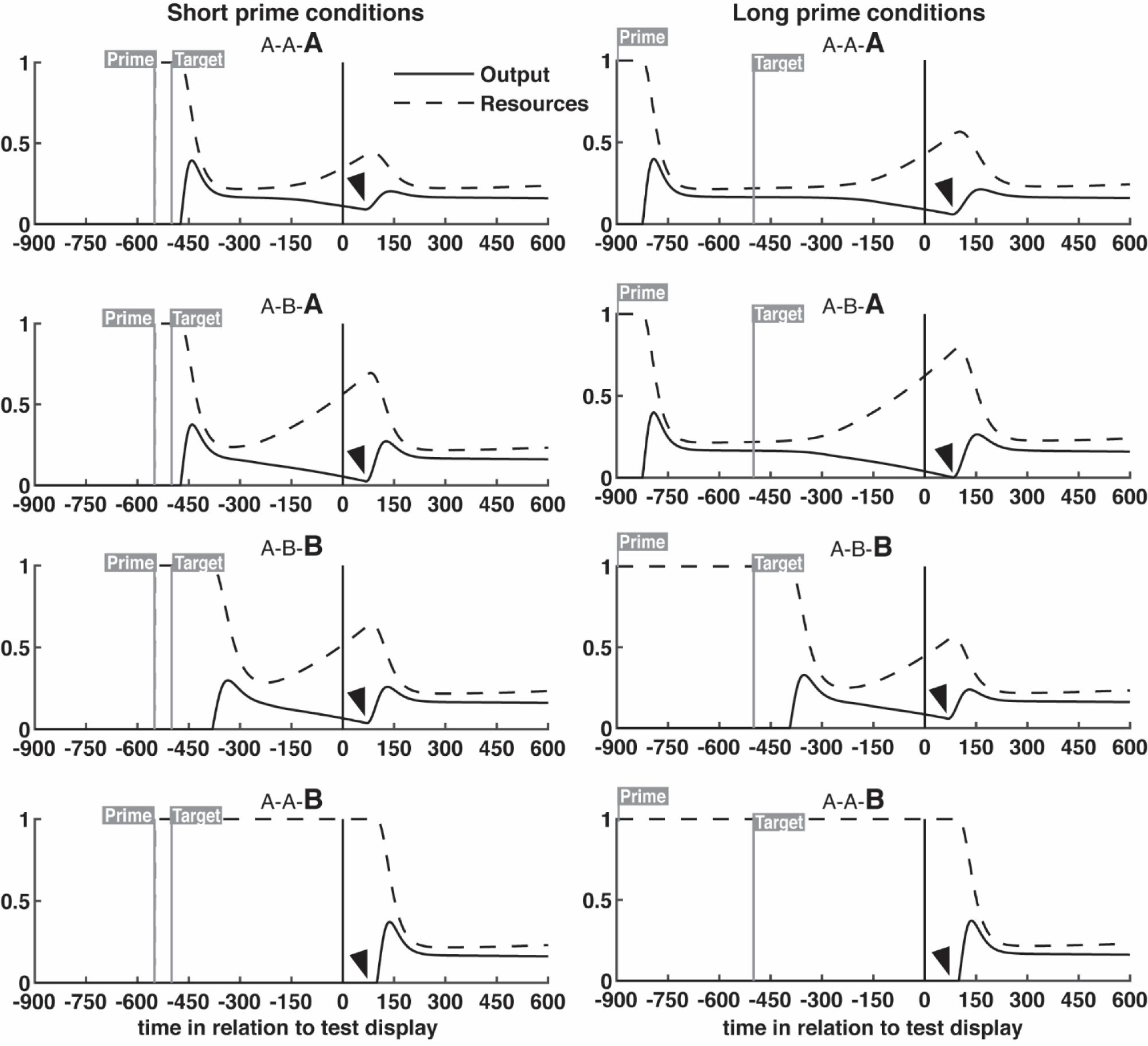
Output and resources for the response word node within the maintained semantic layer of the model. In the model, higher output will consume resources, and decreased resources will reduce output (see “Habituation model” subsection within methods section for details and equations). In this figure, output and resources of the node corresponding to the response word (bolded in each subplot) is shown across a full trial. Residual activations, the lowest output value following response word presentation, are also marked. Lower residual activations cause the model to predict a “different” response, while higher residual activations cause it to predict a “same” response. Accuracy for each of the 8 conditions is determined from the value indicated by the arrow, as compared to a criterion. We consider each condition in turn, working up from the bottom. In the case of the A-A-B conditions, there is no residual activation for B, and so accuracy is good and unchanged by prime duration. In the case of the A-B-B condition, residual activation for B indicates a correct answer of “same”. Residual activate for B increases with increasing prime duration because as habituation for A increases, it does not compete as much with B in the perceptual layers of the model (i.e., a better response to the briefly flashed target word B). In the case of the A-B-A conditions, residual activation from the prime (word A) incorrectly indicates a “same” response. However, for a longer duration prime, word A is habituated, and so there is less residual activation for A, which improves accuracy (enhanced novelty detection). Finally, in the case of A-A-A, residual activation for A correctly indicates a “same” response. However, habituation for word A weakens this residual activation, and reduces accuracy with increasing prime duration (i.e., repetition blindness leads to worse performance).

**Supplementary Fig. 2.**
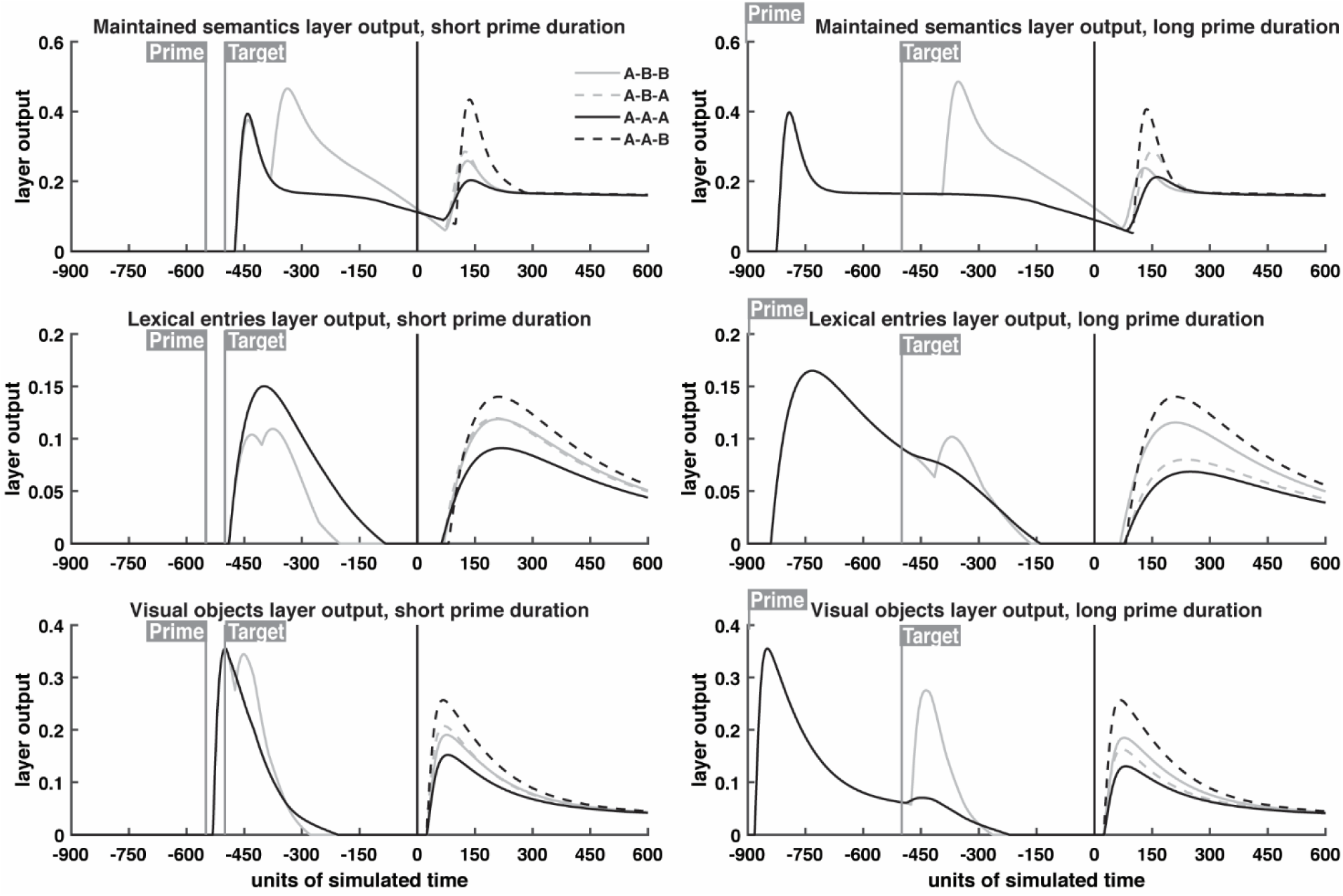
Layer specific activation profiles. The activation profiles of multiple layers were combined to generate N400 and P100/N170 ERP predictions; the above shows the activation of the individual layers. These waveforms were obtained by summing the output of all nodes within a layer at each unit of simulated time. See “Habituation model” subsection within methods section for information on how parameters were obtained.

**Supplementary Fig. 3.**
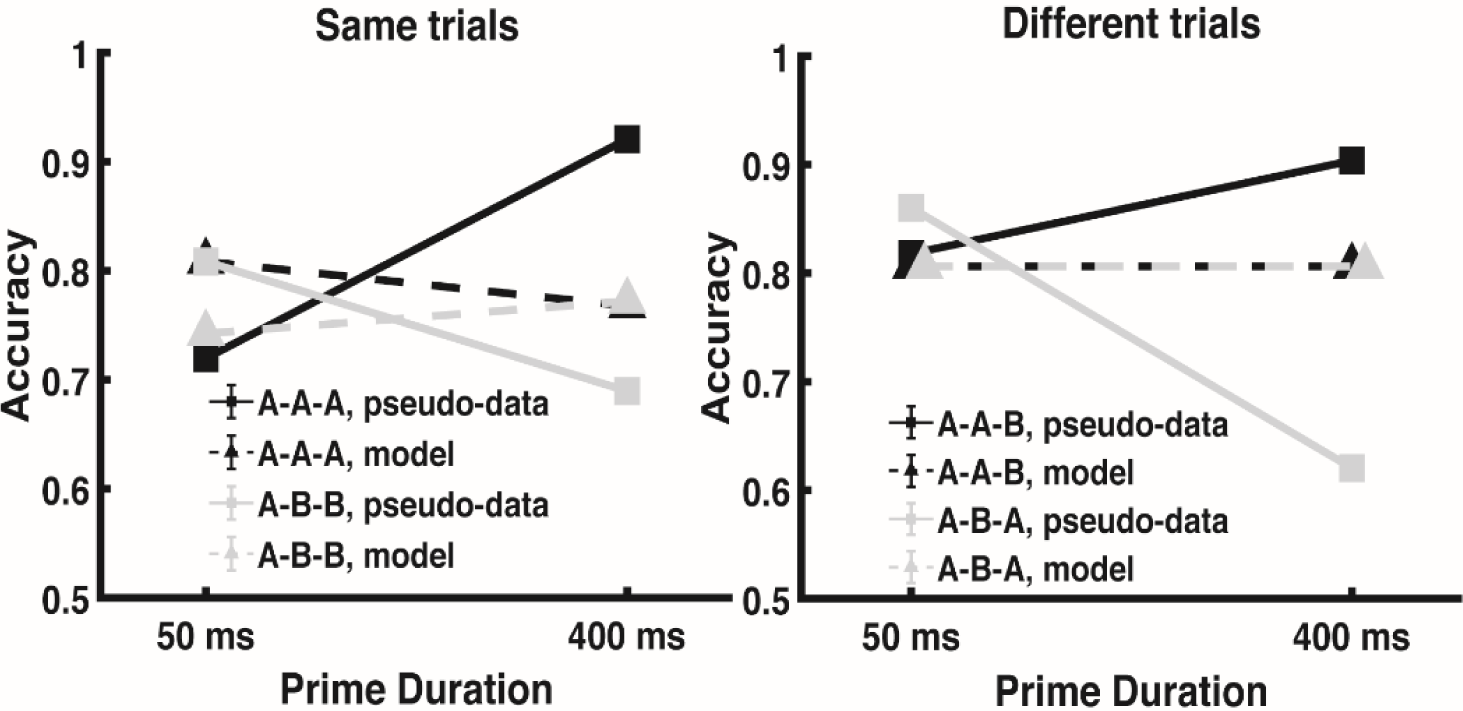
Equivalent model fitting results as compared to the model fits shown in Fig. 4, except that observed data have been time-reversed by labeling the actual 400 ms prime duration conditions as being the 50 ms prime duration conditions and the actual 50 ms prime duration conditions as being the 400 ms prime duration conditions. As in Fig. 4, 5 parameters were optimized in attempt to explain the 8 conditions. Despite having the same 5 free parameters to capture these time-reversed data, the model completely failed to account for the results. The best-fitting χ2 for these time-reversed data was 548.4, which can be contrasted with a χ2 of 40.4 for the results show in Fig. 4. In this case, the best that the model could do was to minimize the role of prime duration to capture the overall average accuracy collapsed across the conditions. This highlights that this is a dynamic systems model, rather than a measurement model. The model is greatly constrained in its predictions for changes across manipulations of duration. More specifically, as prime duration increases, the model necessarily predicts that habituation will increase.

